# Implications of taxonomic and numerical resolution on DNA metabarcoding-based inference of benthic macroinvertebrate responses to river restoration

**DOI:** 10.1101/2021.09.11.459893

**Authors:** Joeselle M. Serrana, Bin Li, Tetsuya Sumi, Yasuhiro Takemon, Kozo Watanabe

## Abstract

Exploring and clearly defining the level of taxonomic identification and quantification approaches for diversity and biomonitoring studies are essential, given its potential influence on the assessment and interpretation of ecological outcomes. In this study, we evaluated the response of benthic macroinvertebrate communities to the restoration or construction of gravel bars conducted in the dam-impacted Trinity River, with the non-dam influenced tributaries serving as the reference sites. We aim to evaluate the performance of different taxonomic and numerical (i.e., abundance vs. presence/absence data) resolutions of DNA metabarcoding with consequent comparison to morphology-based identification and how it affects assessment outcomes. DNA metabarcoding detected 93% of the morphologically identified individuals and provided finer taxonomic resolution. We also detected significant correlations between morphological sample abundance, biomass, and DNA metabarcoding read abundance. We observed a relatively high and significant congruence in macroinvertebrate community structure and composition between different taxonomic and numerical resolutions of both methods, indicating a satisfactory surrogacy between the two approaches and their varying identification levels and data transformation. Additionally, the community-environmental association were significant for all datasets but showed varying significant associations against the physicochemical parameters. Furthermore, both methods identified *Simulium* spp. as significant indicators of the dam-impacted gravel bars. Still, only DNA metabarcoding showed significant false discovery rate proving the method’s robustness compared to morphology-based identification. Our observations imply that coarser taxonomic resolution could be highly advantageous to DNA metabarcoding-based applications in situations where the lack of taxonomic information, e.g., poor reference database, might severely affect the quality of biological assessments.

## INTRODUCTION

Benthic macroinvertebrates are the most commonly used focal groups for the ecological survey of freshwater ecosystems since they serve as valuable indicators of ecosystem health due to their high diversity and different sensitivity to a range of natural and anthropogenic disturbances (Menezes et al., 2010), which have been used to develop biotic indices for extensive monitoring programs (Bush et al., 2019). The accuracy of identification is a key point for macroinvertebrate community-based biomonitoring and assessment. Baird and Hajibabaei (2012) pointed out several constraints that have severely limited the utility of morphology-based macroinvertebrate metrics in broad-scale biomonitoring programs over the last fifty years, e.g., the extensive time required for field sample processing, the identification difficulties for finer taxonomic resolution, the potential identification bias amongst experts, and the general lack of verification of morphology-based identifications.

DNA metabarcoding is a transformative approach to biomonitoring, biodiversity discovery, and ecosystem health assessments in freshwater ecosystems (Cordier et al. 2017; Elbrecht et al. 2017; Serrana et al. 2019), providing solutions to various morphology-based constraints and has been tested and used to study different freshwater habitats and taxonomic groups (e.g., Serrana et al., 2018; Bailet et al., 2019; Ficetola et al., 2020). However, although the method has been reported to identify taxa with finer taxonomic resolution, some argue that this advantage is not necessarily valuable when taxa cannot be linked to a binomial taxonomic name (Bush et al., 2019), which may emerge from incomplete reference DNA libraries (Curry et al., 2018), and mixed template PCR-based challenges, e.g., inadequacy of high performance primers to amplify all relevant taxa (Taberlet et al., 2018).

Taxonomic resolution has substantial effects on ecological study outcomes (Cordier et al., 2017; Pawlowski et al., 2018), but taxonomic details are often set without explicit justification and are usually based on the subjective criteria of sample-processing costs or times (Jones, 2008). Increased taxonomic resolution may increase information and observed variation among communities. This implies that depending on the specificity of the community response to environmental stress, finer taxonomic resolution may either enable better or worse detection of the environmental stress (Bailey, Norris & Reynoldson, 2001). Exploring and clearly defining the community representatives’ taxonomic resolution from diversity and biomonitoring studies is important for understanding the trade-offs associated with different taxonomic levels (Bailey, Norris & Reynoldson, 2001) that is vital for improving the comparability between biogeographically separate programs (Bailet et al., 2019). In particular, the difference in ecological surveys and their employed levels of taxonomic resolution creates potential discrepancies in the results and in the conclusions drawn when comparing the performance of separate programs and procedures (Herbst & Silldorff, 2006). Taxonomic sufficiency – the pragmatic concept of balancing the trade-off between the level of identification against feasibility in ecological studies (Ferrero & Cole, 1992) needs revisiting and discussion, given that species-level identification, which is considered the gold standard in environmental biomonitoring and assessment is not always achievable (Pawlowski et al., 2018), even for DNA metabarcoding-based identifications (Staats et al., 2016; Hleap et al., 2021). Although previous studies have assessed the influence of taxonomic resolution on DNA metabarcoding-based assessments (e.g., Laini et al., 2020), the evaluation of different taxonomic level for identifying taxa without losing significant ecological detail is still largely unexplored.

Furthermore, the numerical resolution of data, i.e., qualitative - presence/absence vs. quantitative – relative or absolute abundance, may influence the outcome of ecological analyses (Mueller, Pander & Geist, 2013; Heino, 2014; Pires et al., 2021). Previous DNA metabarcoding-based macroinvertebrate biomonitoring studies had contrasting reports on numerical resolution. Some suggest the use of presence/absence data (Buchner et al., 2019; Zizka, Geiger & Leese, 2020) since it leads to similar assessment results compared to abundance-based data, while some presented that the quantification of relative species abundance based on read depth information provides better assessment efficiency instead of presence/absence data (Aylagas et al., 2018; Serrana et al., 2019; Meyer et al., 2020).

Sediment deficit downstream reaches due to sediment loading in reservoirs (Petts & Gurnell, 2005) leads to reduced habitat complexity and decreased biodiversity in downstream ecosystems (Graf, 2006). As a resolve, some restoration programs on dam-fragmented rivers employ the construction of gravel bars by gravel augmentation and channel rehabilitation (Ock et al., 2015). Gravel bars provide areas of increased biogeochemical activities due to the enforced hydrodynamic exchange (Sackett et al., 2019), retaining organic matter filtered from surface waters in the hyporheic zone (i.e., the interface between surface and groundwater) for the utilization of river biota, and enhances nutrient cycling with consequent benefits to ecosystem metabolism (Mendoza-Lera & Datry, 2017). Additionally, the wet and dry processes in gravel bars due to water level changes alternately offers terrestrial and aquatic habitats, increasing environmental heterogeneity to provide diverse habitats for different organisms, e.g., fishes (Beechie et al., 2005), microorganisms (Boano et al., 2014), and macroinvertebrates (Merz et al., 2005). Macroinvertebrates have been used as indicators of restoration or enhancement of habitat complexity from gravel bar structures (Lepori et al., 2005; Dunbar et al., 2010) and gravel-bed rivers (Rice & Greenwood, 2001). However, knowledge on the influence of gravel bars on benthic macroinvertebrate communities with specific consideration of the river (i.e., dam fragmented vs. non-dam fragmented river) and reach (gravel bars vs. non-gravel bars) scales are less explored and needs further assessment.

In this study, we aim to evaluate the performance of different taxonomic and numerical resolutions in DNA metabarcoding in comparison to the traditional morphology-based identification and how this would affect the inference of benthic macroinvertebrate responses to river restoration, specifically the monitoring and assessment outcomes on the influence of gravel bar construction or rehabilitation in the dam-impacted Trinity River in California. We hypothesize that different taxonomic levels (i.e., family, genus, species, or haplotype level) and numerical resolution (i.e., absolute abundance or presence/absence data) from both morphology-based and DNA metabarcoding identifications affect the outcome of ecological assessments, i.e., multivariate community pattern, diversity measures, and environmental associations at different spatial scales (i.e., river and reach). Particularly, we expect that the increased information from genus, species, or haplotype-level data relative to a coarser taxonomic level, i.e., family, provides better discrimination in community composition and structure to differentiate test and reference sites at the river (i.e., dam fragmented vs. non-dam fragmented river) and reach (i.e., gravel bars vs. non-gravel bars) scales. We also performed indicator taxa analysis on the datasets of both methods to identify specific genera or species that can be used as indicators of restoration in the gravel bars of the dam-impacted Trinity River in comparison to the reference sites.

## MATERIALS AND METHODS

### Study design and sampling

The Trinity River is a large gravel-bed river located in northwest California, impounded by the Trinity Dam (164 m a.b.l and 3,020 million m^3^ storage) and the smaller Lewiston Dam (28 m a.b.l and 18 million m^3^ storage) located downstream of the Trinity Dam since 1964. The river is under current dam operating guidelines with a mean annual flood of approximately 180 m^3^/s (Gaeuman et al., 2017). The Trinity River Restoration Program, a multi-agency partnership, manages and implements these releases alongside gravel augmentations and mechanical rehabilitations downstream of the Lewiston Dam with the aims of restoring salmonid habitat and dynamic channel processes in the river (USDOI, 2000 as cited in Ock et al., 2015).

The field survey was conducted in the Trinity River along a 60-km river length downstream of the Lewiston Dam in August 2017 to assess the influence of dam-impoundment and gravel bars on benthic macroinvertebrate communities. We performed a paired sampling scheme with two spatial hierarchical scales: i.e., river scale [2 groups: dam-influenced (Trinity River, DAM+) vs. non-dam influenced (tributaries, DAM-)], and reach scale [(2 groups: with gravel bar (GB+) vs. non-gravel bar (GB-)], and their combinations river × reach (4 groups: DAM+GB+, DAM+GB-, DAM-GB+, and DAM-GB-). Six gravel bars and two non-gravel bar reaches of the Trinity River were selected as test sites, while two gravel bars and two non-gravel bar reaches from two non-dam impounded tributaries, i.e., the Rush Creek and the Canyon Creek were sampled as reference sites (**Figure 1**). We sampled each gravel bar site at the bar head (down-welling zone) and bar tail (up-welling zone), while the non-gravel bar segments were sampled at the up- and downstream points approximately 15-m in distance (relative to the average length of the gravel bars). In total, we have 24 sampling points in this study.

**FIGURE 1.**
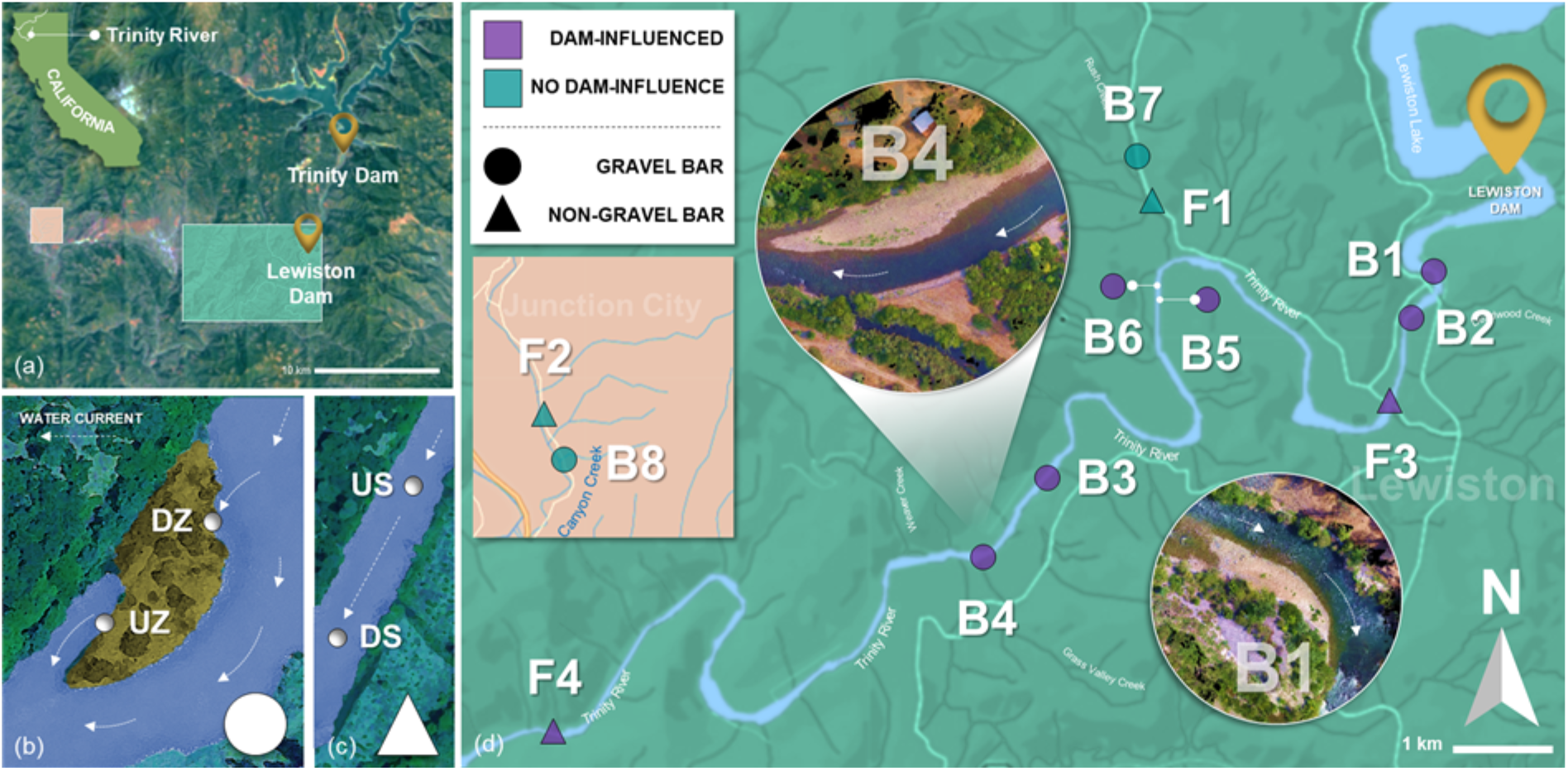
Sampling location. The Trinity River and its tributaries impounded by the Trinity Dam and the Lewiston Dam in California, USA (a). Gravel bars were sampled at the head (down-welling zone; DZ) and tail (up-welling zone; UZ) points (b). Non-gravel bar sites were assessed, and samples were collected on up-(US) and downstream (DS) points (c). In total, eight gravel bars and four non-gravel bar sites were assessed in the study; gravel bars B1 and B4 drone photos shown as examples (d). Parts of the map was generated from Google Earth Pro (version 7.3.2.5776; https://www.google.com/earth/).

Physico-chemical characteristics were measured from all sampling points (**Table S1**). Water samples for water quality analysis were collected following the procedures instructed by the United States Environmental Protection Agency (USEPA), kept in an icebox, and sent to the PHYSIS Environmental Laboratories, Inc. (Anaheim, CA, USA) for water quality analysis, i.e., ammonium nitrogen (NH_4_-N), nitrate nitrogen (NO_3_-N), ash-free dry mass (AFDM), and total suspended solids (TSS). pH and electric conductivity (EC) was measured via the LAQUAtwin COND (Model Y071L, Horiba) and pH meters (Model Y017, Horiba). Water temperature and dissolved oxygen were measured with a DO meter (Model OM-51, Horiba, Ltd., Kyoto, Japan).

Benthic macroinvertebrates were collected using a Surber net (30 cm × 30 cm, mesh size 500 μm) at three random locations to cover most habitats (e.g., riffles and pools) at each sampling point. The Surber replicates were pooled together and directly preserved in 99% ethanol in the field. In the laboratory, the collected macroinvertebrates were sorted from debris, e.g., substrate and non-target organic matters, and morphologically identified at the lowest taxonomic level when possible (usually genus or family) under a stereomicroscope using the taxonomic keys for the identification of aquatic insects in North America (Merritt, Cummins & Berg, 2008). Sorted and morphologically identified samples were then kept in new ethanol and stored at 4°C before processing for molecular analysis.

### DNA extraction, amplicon library construction, and sequencing

The morphologically identified and sorted samples from each point were dried overnight at room temperature in sterile Petri dishes to remove the remaining ethanol. The dry biomass of each taxon was measured on a Sartorius RC 210D semi-microbalance. Individuals from each taxon were then pooled as a bulk sample and homogenized by grinding in several 2-mL tubes. Genomic DNA was extracted using the DNeasy Blood & Tissue Kit (Qiagen, Inc.) following manufacturer instructions. DNA concentrations were quantified with the QuantiFluor dsDNA system (Promega, Madison, WI, USA) on the Quantus Fluorometer (Promega, Madison, WI, USA) and normalized to 50 ng/μL DNA for PCR.

A two-step PCR protocol following the procedures of Elbrecht and Steinke (2019) was applied for PCR amplification and tagging. For the first PCR step, we amplified the target fragment using the BF2+BR2 primer set designed explicitly for benthic macroinvertebrates and evaluated using both mock and kick-net samples (Elbrecht & Leese, 2017). The PCR master mix consists of 0.25 μl Phusion, 0.75 μl Dimethyl sulfoxide (DMSO), 1 μl dNTPs, 1.25 μl each of the forward and reverse primers (10 μM), five μl HF Buffer (New England Biolabs), and 15.5 μl of PCR-grade water. PCR cycling conditions were 30 seconds of initial denaturation at 98°C, followed by 25 cycles of 10 seconds denaturation at 98°C, 30 seconds annealing at 55°C, 30 seconds extension at 72 °C, and a final extension step of 5 minutes at 72°C. For the second PCR, we used fusion primers, which include inline tags and Illumina sequencing tails. One μl of the first PCR was used as the template. PCR master mix and conditions are the same as the first PCR step, but the cycles were adjusted to 15×. The second PCR products were purified by QIAquick PCR Purification Kit (QIAGEN, Inc., Valencia, CA), and the DNA concentration was quantified with qPCR via the KAPPA Illumina Library qPCR Quantification kit (Kappa Biosystems, Wilmington, MA, USA) and qualified with the High-Sensitivity DNA chip (Agilent BioAnalyzer, Palo Alto, CA, USA) for amplicon library assessment. PCR water was used as a negative control to monitor contamination from DNA extraction and library construction to post-amplification library quantity and quality verification, and no quantifiable amplicon was detected for further analysis. The 72 amplicon libraries of the 24 samples with triplicates were then normalized to 4 nM and pooled. Finally, 600 μl of a 6 pM denatured pooled library with PhiX (final concentration 15%; Illumina, Inc.) was prepared, and a 300-bp paired-end sequencing was performed using the MiSeq Reagent Kit v3 (Illumina, Inc.).

The generated raw sequence data were deposited into the National Center for Biotechnology Information (NCBI) Sequence Read Archive (SRA) under the accession number SRR12620160.

### Bioinformatics and data processing

The raw Illumina paired-end reads were demultiplexed according to sample tags via the R package JAMP v.0.67 (http://github.com/VascoElbrecht/JAMP) (Elbrecht et al., 2018) and were quality-checked with FastQC (https://www.bioinformatics.babraham.ac.uk/projects/fastqc/). The primer sequences were trimmed using Cutadapt v.2.1 (Martin, 2011), and the subsequent read processing steps, i.e., quality filtering, merging, and inference of amplicon sequence variants (ASVs) were performed following the denoising pipeline of the DADA2 v.1.12 package (Callahan et al., 2016) in R v.3.6.2 (R Core Team, 2019). The forward and reverse reads were trimmed at a minimum length of 220 and quality filtered using a maximum expected error (-maxee) of 3 and 5. The remaining sequences were denoised, and the forward and reverse sequences were merged. ASVs were inferred from the sequence data while subsequently removing chimeric sequences and singletons.

For the taxonomic assignment, the ASV sequences were matched to the Barcode of Life Database (BOLD, Barcode of Life Data System, http://www.boldsystems.org/; accessed at September 1, 2020) using the python program BOLDigger (https://github.com/DominikBuchner/BOLDigger) (Buchner & Leese, 2020). The best-fitting hit option from the JAMP pipeline was performed, wherein different thresholds, i.e., 98% for species level, 95% for the genus, 90% for family, 85% for order level, and <85% for class level, were used as a best-fitting hit parameter. An interactive Krona chart was used to visualize the total individual count and absolute read abundances within the complex hierarchies of the taxonomic assignments (Ondov et al., 2011).

### Statistical analysis

Pearson’s pairwise correlation test was performed to assess the relationship between morphologically-identified sample abundance and biomass per taxa against the number of reads in the metabarcoding data. The shared and unique taxonomic assignment and ASVs at the river × reach scale was visualized with UpSetR plots (Lex et al., 2014). The read counts per ASV and sample data, along with the taxonomic identifications and sample descriptors were merged into phyloseq objects using the phyloseq v.1.32.0 package (McMurdie and Holmes 2011). For the subsequent analysis, the DNA metabarcoding reads and the morphologically-identified individual counts in each sample were normalized using median depth. Median normalization was performed as recommended in Pereira et al. (2018) since it provides a robust alternative to total count that is less affected by highly abundant samples. Four datasets with different taxonomic resolutions were created from the DNA metabarcoding dataset, i.e., family, genus, species, and ASV-level, while there were the family and genus-level for the morphological dataset. Absolute abundance and presence/absence datasets were also created from these four DNA metabarcoding and two morphological datasets.

Alpha diversity metrics, i.e., Chao1 richness, Shannon diversity, Pielou’s J evenness, Berger-Parker dominance, and the rare abundance index were calculated and visualized to identify the changes in community structure between each scale using the *plot_alpha_diversities* function (microbiomeutilities; Shetty & Lahti, 2018). Statistical differences of the alpha diversity metrics among each scale were tested using ANOVA and pairwise comparisons via multiple t-tests. Pearson correlation coefficients between the alpha diversity metrics of each taxonomic level and methods were calculated using the *rcorr* function in the Hmisc package. Additionally, a meta-regression analysis was performed to explore the simultaneous effects of all seven environmental variables (log-transformed) with the richness estimate (i.e., Chao1) as the dependent variable using the *lm* function of the stats package. The *plot_models* function of the sjPlot package was then used to plot and compare regression coefficients with confidence intervals of multiple regression models in one plot.

To observe the spatial differences between the communities, beta diversity was estimated based on Bray-Curtis distances and assayed by non-metric multidimensional scaling (NMDS) using the *plot_ordination* function from the phyloseq package. Permutational multivariate analysis of variance (PERMANOVA) (vegan; Oksanen et al., 2013) was performed among the scales to test the statistical significance of the between-group distances based on the NMDS ordination via the *adonis2* function in the vegan package. Procrustes tests were employed via the *protest* function with 9,999 permutations to test if different taxonomic or numeric resolutions provided similar community structure. A canonical correspondence analysis (CCA) was performed to visualize and determine the environmental variables responsible for explaining community composition variation amongst sampling points with significance. Before the analysis, the seven environmental variables were evaluated for multi-collinearity via variance inflation factors (VIF) for constraining parameters (VIF <5) using the usdm package (Naimi 2015). Additionally, a homogeneity of multivariate dispersion (PERMDISP) analysis was employed to assess if differences in heterogeneity in environmental parameters existed among the samples based on the river, reach, and river × reach scales.

Indicator taxa analysis was performed to the genus-level morphology and metabarcoding datasets to identify and compare the indicator taxa between the two methods based on the river × reach category using the *multipatt* function implemented in the indicspecies package in R with 9,999 permutations (De Caceres et al., 2016). The associations were further assessed using a false discovery rate estimation by adjusting the *p*-values for multiple testing using the Benjamini-Hochberg procedure (*p*.*adjust* function in R) (Strimmer, 2008). Furthermore, the linear discriminant analysis effect size (LEfSe) test was conducted using the Galaxy implementation of LEfSe (http://huttenhower.org/galaxy) (Segata et al., 2011) [parameters: *p* < 0.05, *q* < 0.05, LDA > 2.0, multi-class analysis strategy set to one-against-all (less strict)] to identify which indicator taxa significantly explained differences in community composition between the river × reach category. All the data visualization and statistical analyses were performed in R v.3.6.2 (R Core Team, 2019).

## RESULTS

### Taxonomic identification based on morphology and DNA metabarcoding

We collected a total of 4,053 macroinvertebrates and morphologically identified 39 multi-level taxa from 2 phyla, 3 classes, 9 orders, 31 families, 25 genera, and 2 species. Almost all samples had family level assignments (31 families), except for the individuals identified as order Haplotaxida (Oligochaeta). Only 68% (2,737 individuals) were assigned at the genus level, from which 99 and 37 individuals had species-level assignments, i.e., *Calineuria californica* and *Pteronarcys californica*. A total of 7,399,885 paired-end reads were demultiplexed from the raw sequence data. After read processing, i.e., quality filtering, denoising, and paired-end merging, 4,098,620 non-chimeric reads (∼421-bp) were retained and assigned to 1,210 ASVs. From this, 4,083,001 reads from 1,112 ASVs were assigned to macroinvertebrate taxa. ASVs assigned as bacteria, fungi, and plants (17 ASVs; 11,703 reads) and ASVs without taxonomic identifications (79 ASVs; 3,916 reads) were discarded from the dataset. Out of the total taxonomically assigned reads, 91% (3,701,486 reads) had a family level assignment, while 87% (3,561,696 reads) and 65% (2,640,658) had genera and species level assignment. The sample, read abundance, and ASV count were summarized in detail in **Table 1**. Interactive Krona charts were provided as an additional file to visualize the total individual count and absolute read abundances of the taxonomic assignments from the two methods.

**TABLE 1.**
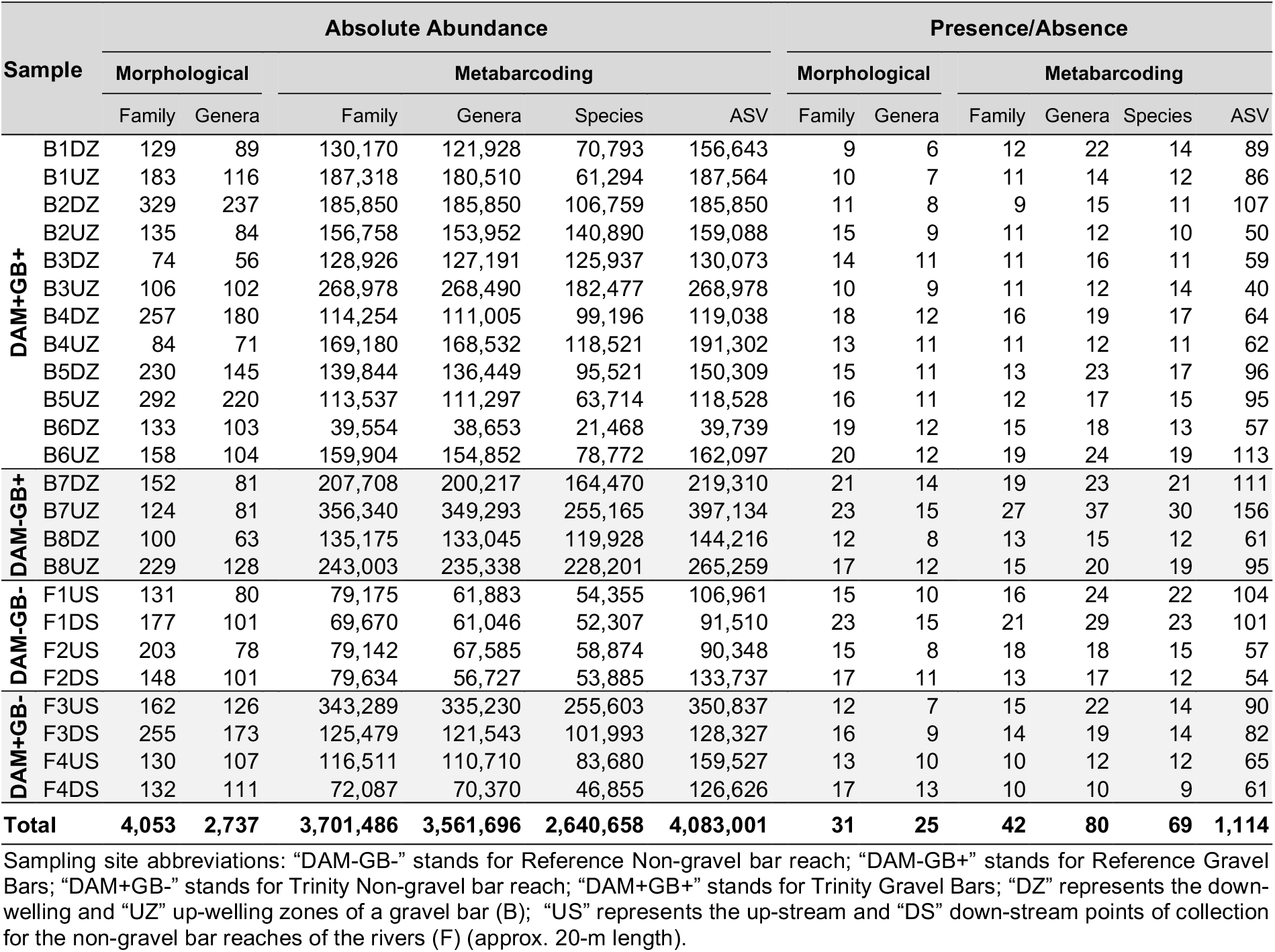
Absolute abundance and presence/absence count of morphologically identified samples and DNA metabarcoding reads.

From the morphology-based identification, the most abundant order was Diptera (35%), followed by the EPT orders: Ephemeroptera (33%), Plecoptera (17%), and Trichoptera (11%). However, DNA metabarcoding detected Plecoptera (39%) as the order with the most reads, followed by Ephemeroptera (33%), Trichoptera (11%), and Diptera (8%). See **Figure S1** for the relative abundance of order-level assignment of the two identification methods per sampling site. DNA metabarcoding detected 93% (3,767 individuals) of the morphologically identified taxa, accounting for 83% of the reads (**Figure 2a** to **2c**). Only 7% (286 individuals) were false negatives, representing three genera, i.e., *Mataeopsephus* (Coleoptera), *Nemoura* (Plecoptera), and *Rhitrogena* (Ephemeroptera). The reads related to these genera were possibly assigned to coarser taxonomic levels, given that some ASVs have unclassified class, order, and family identifications. The remaining 17% (696,709 reads) consists of 44 taxa, i.e., 13 species, 12 genera, and 18 unclassified ranks unique to the DNA metabarcoding dataset which are considered false positive detections. We found significant positive linear correlations between sample abundance, sample biomass, and read abundance among the 36 taxa commonly found in the morphological and DNA metabarcoding datasets presented in **Figure 2d** to **2f**.

**FIGURE 2.**
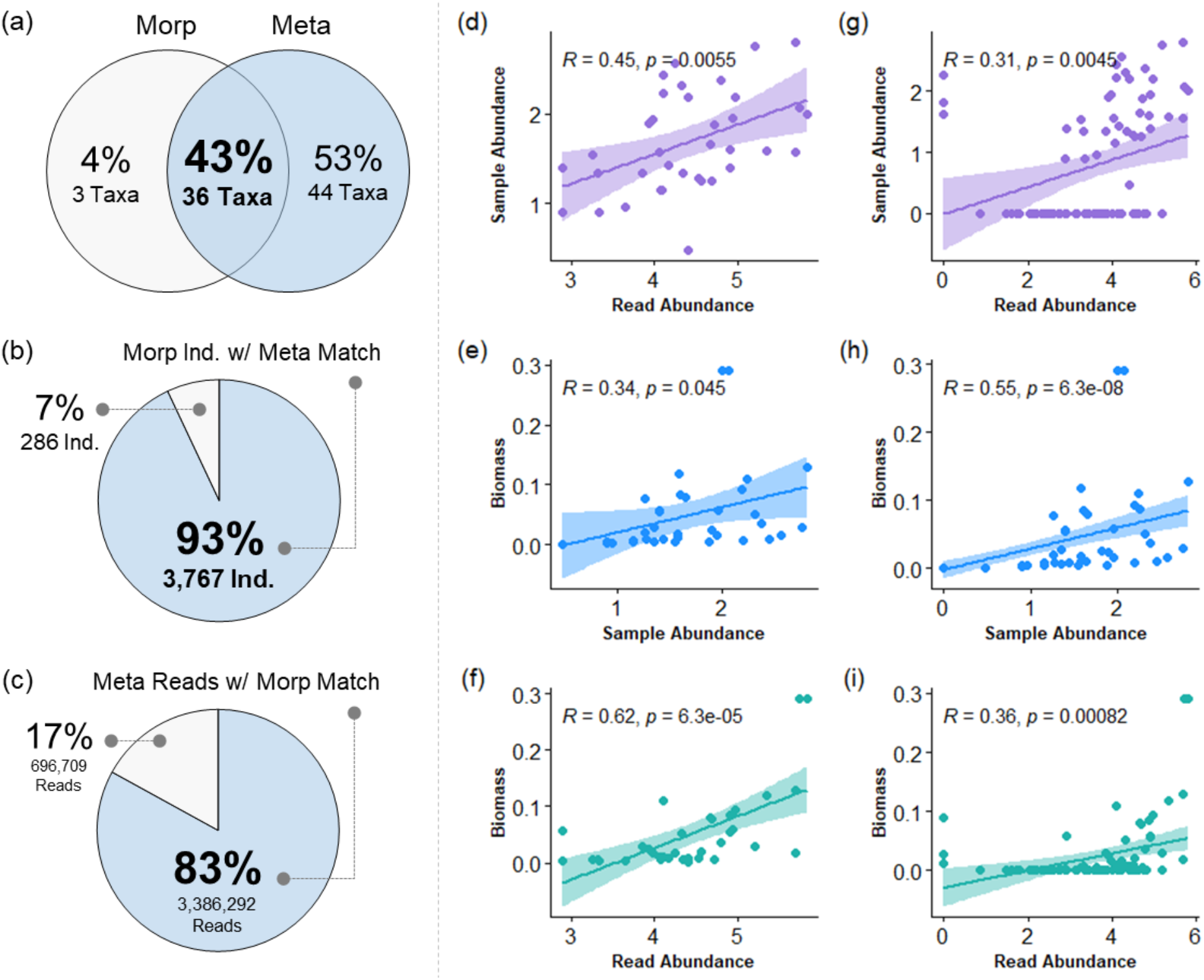
Venn diagram showing the shared and unique taxa between the morphological (“Morp”) and metabarcoding (“Meta”) methods (a). The pie charts present the number of morphologically identified individuals (b) and their corresponding read abundance (c) detected in metabarcoding. Pearson correlations between the sample abundance (i.e., individual count), biomass (i.e., dry biomass in grams), and metabarcoding read abundance (i.e., absolute sequence count). Showing the analysis excluding (d-f), and including (g-i) false positive detections. Data were log-transformed prior to analysis.

### Richness and diversity measures

The shared and unique taxa at the river × reach category for the different taxonomic resolution datasets of both methods were presented in **Figure 3**. Thirteen morphologically identified families and fourteen DNA metabarcoding families were shared by all the river × reach groups, i.e., DAM+GB+, DAM+GB-, DAM-GB+, and DAM-GB-. Both methods identified fourteen genera present in all groups, while 20 species were shared between the groups from the DNA metabarcoding dataset. The pairwise correlation of the alpha diversity estimates of the morphological and DNA metabarcoding datasets at different taxonomic resolutions was presented in **Figure 4**. The pairwise correlations between the datasets with the different taxonomic and numerical resolutions (absolute abundance or presence/absence) were significantly positive in 47 pairs (taxonomic: 20; numerical: 27) for Chao1 richness, 42 pairs (taxonomic: 18; numerical: 24) for Shannon diversity, 18 pairs (taxonomic: 12; numerical: 6) for Berger-Parker dominance index, and 25 pairs (taxonomic: 11; numerical: 14) for rare taxa abundance index out of 66 pairs. We observed varying statistical differences of the alpha diversity metrics among the scales (i.e., river, reach, or river × reach) at different taxonomic and numerical resolutions presented in **Table S1**. Remarkably, we observed no significant difference in all the datasets for the mean alpha diversity values at the reach scale. Another notable observation was that only the family and species-level absolute abundance DNA metabarcoding datasets showed a significant difference between the river × reach scale.

**FIGURE 3.**
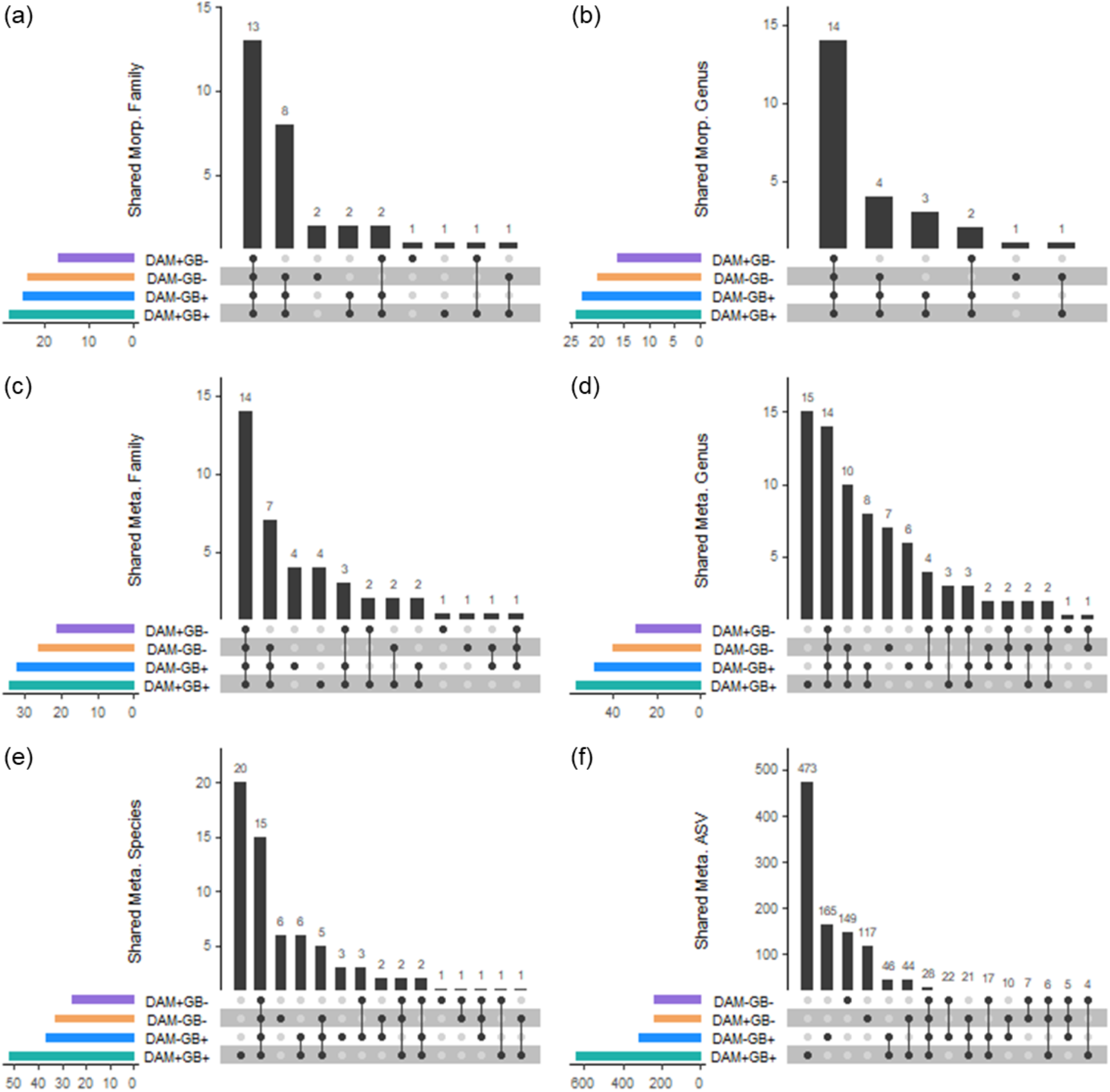
Number of taxa shared between River × Reach scale for the morphology-based (a-b) and metabarcoding (c-f) data.

**FIGURE 4.**
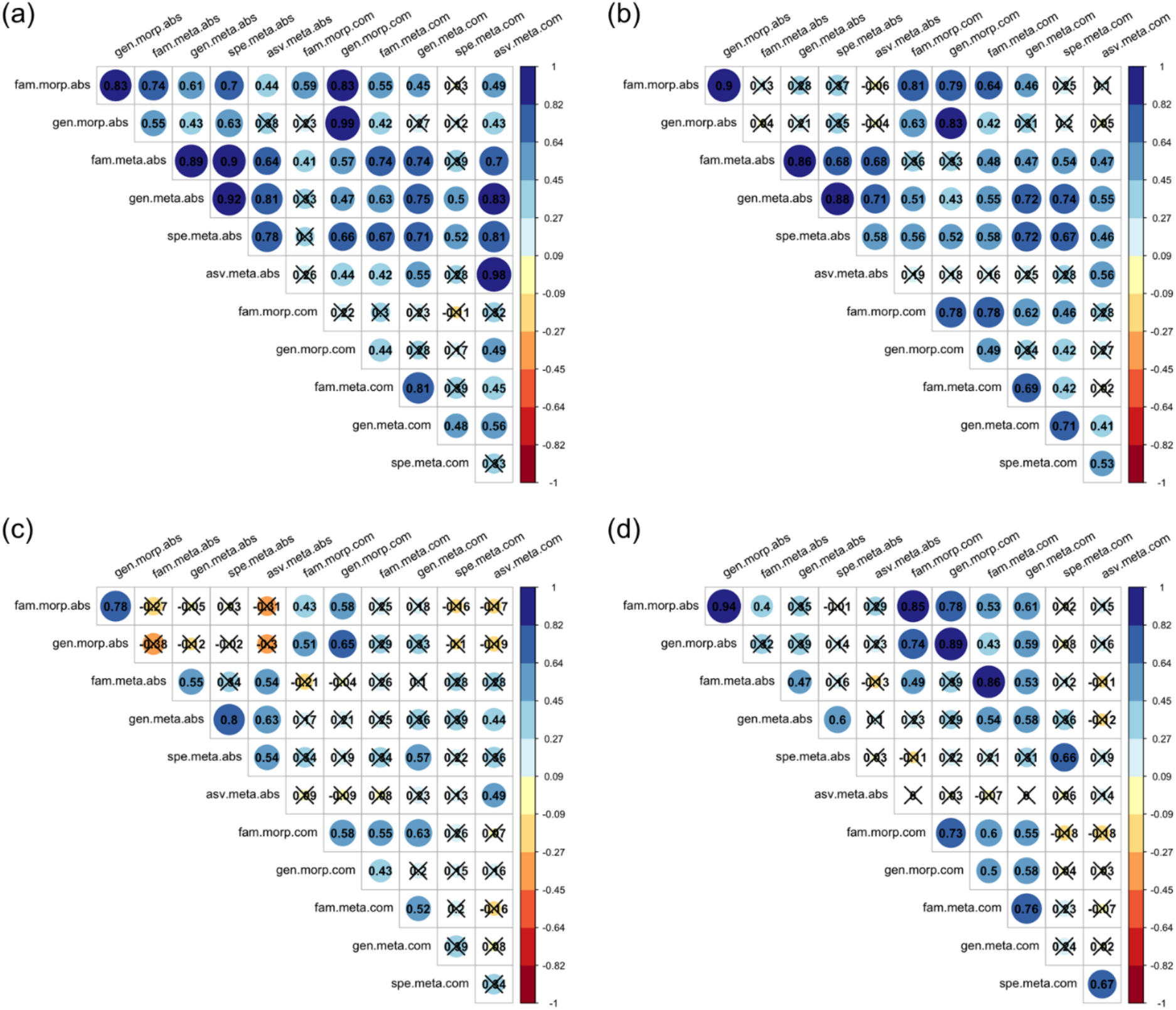
Pairwise Pearson’s correlation of alpha diversity estimates. Chao1 richness (a), Shannon diversity (b), Berger-Parker dominance index (c), and rare taxa abundance (d). Correlation boxes with Xs are not significant at *p* = 0.05. “morp” indicates morphologically identified; “meta” for DNA metabarcoding; “gen” for genus-level identification; “fam” for family-level; “spe” for species–level; “asv” for the amplicon sequence variant (ASV) level dataset.

**FIGURE 5.**
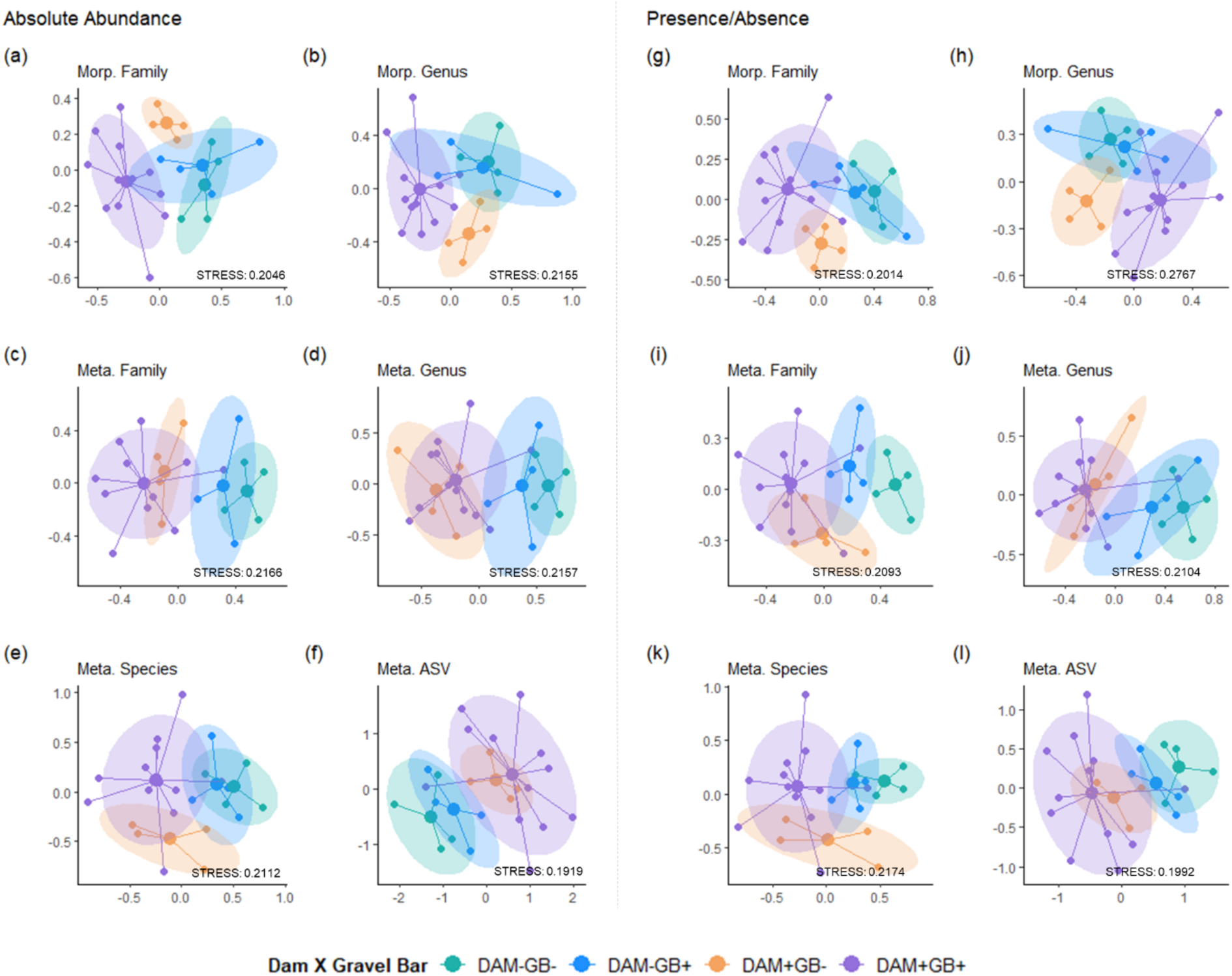
Non-metric multidimensional scaling (NMDS) plot on Axes 1 (x-axis) and 2 (y-axis) based on Bray-Curtis dissimilarity for absolute abundance (a-f) and presence/absence (g-l) data of the macroinvertebrates detected by morphological (“Morp”) and DNA metabarcoding-based (“Meta”) methods.

### Community structure and composition, and environmental relationship

To assess whether different taxonomic and numerical resolution interferes in the homogeneity of multivariate dispersion within each scale, the community structure and composition of the identified benthic macroinvertebrates were assessed with NMDS based on Bray-Curtis distance. Ordination plots were relatively similar for all the taxonomic levels, and numerical resolution on both methods, wherein the communities of the samples from the dam-impacted sites (from the Trinity River) with and without gravel bars were closely clustered in the ordination space, and the non-dam impacted samples clustered together (**Figure 4**). The visual similarity between the NMDS ordinations between both methods’ different taxonomic and numerical datasets was further supported by the Procrustes analyses, which revealed significant positive correlations (**Figure S2**). However, relatively few comparisons had high correlation coefficient values (from 0.90 to 0.98).

The PERMANOVA analyses revealed that community structure and composition at the river scale, dam-influenced rivers, significantly differed against the non-dam influenced rivers (PERMANOVA, *p* = 0.001) for all the different taxonomic and numerical resolution datasets of both methods (**Table 2**). Almost all datasets showed a significant difference between the communities at the reach scale (PERMANOVA, *p* < 0.05), except for the ASV-level datasets and the family-level morphology-based presence/absence dataset. These results indicate the strong influence of dams and gravel bars on the benthic macroinvertebrates’ community structure. However, no significant difference between the community composition at the river × reach scale was found.

**TABLE 2.** Permutational analysis of variance (PERMANOVA) results for each scale.

| Taxonomic Rank |  |  | River |  |  | Reach |  |  | River × Reach |  |  |
| --- | --- | --- | --- | --- | --- | --- | --- | --- | --- | --- | --- |
|  |  |  | *F.Model* | *R*<sup>2</sup> | *Pr(>F)* | *F.Model* | *R*<sup>2</sup> | *Pr(>F)* | *F.Model* | *R*<sup>2</sup> | *Pr(>F)* |
| Absolute Abundance | Morp | Family | 6.15 | 0.21 | **0.001** | 2.21 | 0.07 | 0.038 | 1.46 | 0.05 | 0.171 |
|  |  | Genus | 4.44 | 0.16 | **0.001** | 2.42 | 0.09 | 0.015 | 1.22 | 0.04 | 0.282 |
|  | Meta | Family | 6.15 | 0.21 | **0.001** | 2.25 | 0.08 | 0.031 | 1.32 | 0.04 | 0.241 |
|  |  | Genus | 5.65 | 0.20 | **0.001** | 2.16 | 0.07 | 0.027 | 1.01 | 0.04 | 0.398 |
|  |  | Species | 3.95 | 0.15 | **0.001** | 2.02 | 0.07 | 0.022 | 1.21 | 0.04 | 0.284 |
|  |  | ASV | 2.51 | 0.10 | **0.001** | 1.19 | 0.05 | 0.142 | 1.23 | 0.05 | 0.122 |
| Presence/Absence | Morp | Family | 7.25 | 0.24 | **0.001** | 1.97 | 0.06 | 0.074 | 1.39 | 0.05 | 0.239 |
|  |  | Genus | 3.15 | 0.12 | **0.008** | 2.31 | 0.09 | 0.035 | 1.35 | 0.05 | 0.276 |
|  | Meta | Family | 7.31 | 0.23 | **0.001** | 2.87 | 0.09 | 0.016 | 1.03 | 0.03 | 0.442 |
|  |  | Genus | 6.18 | 0.21 | **0.001** | 2.73 | 0.09 | 0.011 | 0.84 | 0.03 | 0.579 |
|  |  | Species | 4.65 | 0.16 | **0.001** | 2.54 | 0.09 | 0.014 | 1.18 | 0.04 | 0.319 |
|  |  | ASV | 2.47 | 0.10 | **0.001** | 1.23 | 0.05 | 0.101 | 1.20 | 0.05 | 0.148 |

Significant values are indicated in highlight (<0.05) and bold print (<0.01). "Morp" indicates morphologically identified; "Meta" for DNA metabarcoding.

Environmental and water quality parameters were assessed for all twenty-four sampling sites (**Table S2**). **Figure S3a** to **S3c** presents the distribution of each environmental variable per scale. However, a PERMDISP2 analysis (**Table S3**) only revealed two of the seven environmental variables significantly different amongst the scales, i.e., electric conductivity (EC) significantly different for both river and river × reach, and total suspended solids (TSS) significant different for reach and river × reach. The variation in the correlations of the different datasets in relation to the seven environmental variables is also shown in **Figures S3d** and **S3e**. The CCA analysis models to test the influence of environmental variables on community composition were significant for all the different taxonomic and numerical datasets of both methods, with *R*^*2*^ values ranging from 0.32 to 0.46. **Figure S4** presents the CCA biplots. However, the different datasets showed a difference in significant associations against the seven physicochemical parameters tested (**Table 3**). For example, pH significantly influenced the benthic macroinvertebrate community composition for the metabarcoding family (presence/absence only), species, and ASV-level datasets. EC was significant for the morphology-based family and metabarcoding family and genus-level datasets. Notably, total suspended solids (TSS) were only significant at the morphology-based family-level absolute abundance dataset.

**TABLE 3.**
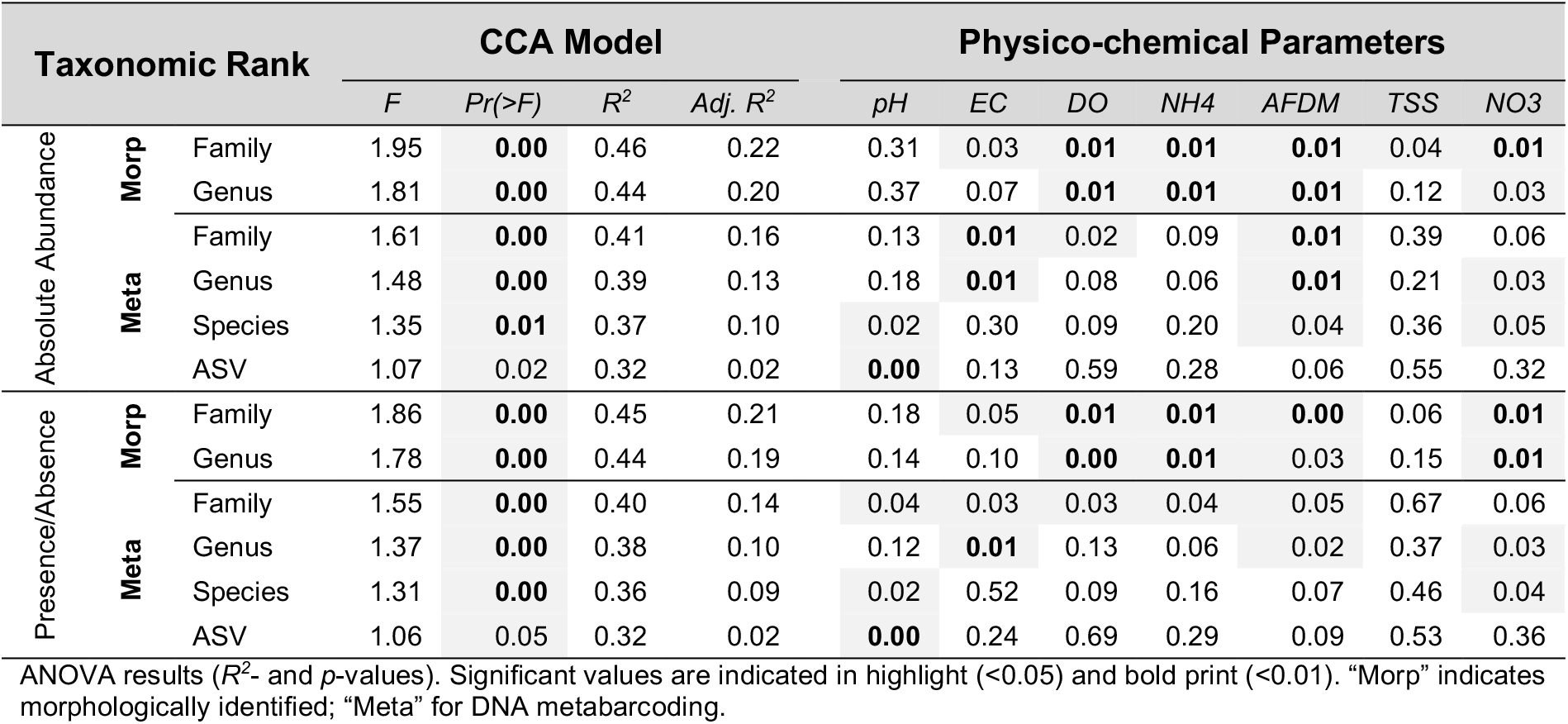
Canonical correspondence analysis (CCA) of the environmental variables.

| Taxonomic Rank |  |  | CCA Model |  |  |  | Physico-chemical Parameters |  |  |  |  |  |  |
| --- | --- | --- | --- | --- | --- | --- | --- | --- | --- | --- | --- | --- | --- |
|  |  |  | *F* | *Pr(>F)* | *R*<sup>2</sup> | *Adj. R*<sup>2</sup> | *pH* | *EC* | *DO* | *NH4* | *AFDM* | *TSS* | *NO3* |
| Absolute Abundance | Morp | Family | 1.95 | **0.00** | 0.46 | 0.22 | 0.31 | 0.03 | **0.01** | **0.01** | **0.01** | 0.04 | **0.01** |
|  |  | Genus | 1.81 | **0.00** | 0.44 | 0.20 | 0.37 | 0.07 | **0.01** | **0.01** | **0.01** | 0.12 | 0.03 |
|  | Meta | Family | 1.61 | **0.00** | 0.41 | 0.16 | 0.13 | **0.01** | 0.02 | 0.09 | **0.01** | 0.39 | 0.06 |
|  |  | Genus | 1.48 | **0.00** | 0.39 | 0.13 | 0.18 | **0.01** | 0.08 | 0.06 | **0.01** | 0.21 | 0.03 |
|  |  | Species | 1.35 | **0.01** | 0.37 | 0.10 | 0.02 | 0.30 | 0.09 | 0.20 | 0.04 | 0.36 | 0.05 |
|  |  | ASV | 1.07 | 0.02 | 0.32 | 0.02 | **0.00** | 0.13 | 0.59 | 0.28 | 0.06 | 0.55 | 0.32 |
| Presence/Absence | Morp | Family | 1.86 | **0.00** | 0.45 | 0.21 | 0.18 | 0.05 | **0.01** | **0.01** | **0.00** | 0.06 | **0.01** |
|  |  | Genus | 1.78 | **0.00** | 0.44 | 0.19 | 0.14 | 0.10 | **0.00** | **0.01** | 0.03 | 0.15 | **0.01** |
|  | Meta | Family | 1.55 | **0.00** | 0.40 | 0.14 | 0.04 | 0.03 | 0.03 | 0.04 | 0.05 | 0.67 | 0.06 |
|  |  | Genus | 1.37 | **0.00** | 0.38 | 0.10 | 0.12 | **0.01** | 0.13 | 0.06 | 0.02 | 0.37 | 0.03 |
|  |  | Species | 1.31 | **0.00** | 0.36 | 0.09 | 0.02 | 0.52 | 0.09 | 0.16 | 0.07 | 0.46 | 0.04 |
|  |  | ASV | 1.06 | 0.05 | 0.32 | 0.02 | **0.00** | 0.24 | 0.69 | 0.29 | 0.09 | 0.53 | 0.36 |

ANOVA results (*R*<sup>2</sup>- and *p*-values). Significant values are indicated in highlight (<0.05) and bold print (<0.01). "Morp" indicates morphologically identified; "Meta" for DNA metabarcoding.

### Indicator taxa analysis

We compared the indicator genera identified between the morphology-based and DNA metabarcoding methods based on the genus-level datasets. Three genera, i.e., *Rhyacophila, Simulium*, and *Nemoura* were identified as indicators with a *p*-value significant at 0.05 from the morphology-based dataset, while thirteen were identified from the DNA metabarcoding dataset, with *Wormaldia, Hydropsyche, Antocha, Attenella, Haemaphylis, Ordobrevia*, and *Simulium* having significant *p*-values at 0.05, and false discovery rate adjusted *p*-values less than 0.10. See **Table S4** for the group associations of the aforementioned indicator taxa at the river × reach scale. Only *Simulium* (Diptera) was reported for both methods, notably an indicator of the dam-impacted Trinity River’s gravel bar sites (DAM+GB+).

Moreover, an additional LEfSe analysis was performed at the multi-taxa dataset (i.e., all ASVs with varying taxonomic assignment) of both methods. Thirteen and 59 significantly discriminative features (i.e., taxa) with an LDA score of > 2.0 out of the 13 and 105 before internal Wilcoxon were selected from the morphology-based and DNA metabarcoding datasets. See Supplementary **Figures S5** and **S6** for the taxa identified that significantly explained differences in community compositions between the river × reach scales. Notably, LEfSe analyses identified genus *Simulium* as an indicator of the dam-impacted Trinity River’s gravel bar sites (DAM+GB+) for both methods. However, DNA metabarcoding provided species-level assignments and identified *S. vittatum* alongside *Dicosmoecus gilvipes, Hesperoperla pacifica*, and *Sialis californica* as indicator species of the dam-impacted rivers with gravel bars (DAM+GB+).

## DISCUSSION

The main objective of this study is to evaluate the influence of the construction or rehabilitation of gravel bars in the dam-impacted Trinity River at the river (i.e., dam fragmented vs. non-dam fragmented river) and reach (i.e., gravel bars vs. non-gravel bars) scales by performing DNA metabarcoding and morphology-based identifications of benthic macroinvertebrate communities of different taxonomic levels, i.e., family, genus, species, and haplotype (ASV-level), and numerical resolutions, i.e., absolute abundance or presence/absence data.

### Congruence between DNA metabarcoding and morphology-based identifications

We confirm the effectiveness of DNA metabarcoding as an alternative method to overcome certain limitations of the conventional morphology-based identification. DNA metabarcoding detected most of the morphologically identified taxa and provided finer taxonomic resolution, up to species and haplotype-level information. This advantage was previously demonstrated in macroinvertebrate taxa studies (e.g., Baird & Hajibabaei, 2012; Elbrecht et al., 2017; Emilson et al., 2017; Serrana et al., 2019). The false positive detection between the two methods could be due to the lack of reference sequences and misclassifications in the database or primer bias effects (Carew et al., 2017; Laini et al., 2020). Regardless, we report a significant correlation between morphological abundance and sequence read abundance, between morphological abundance and biomass, and between sequence read abundance and biomass. These results were consistent with previous DNA metabarcoding studies, which reported a positive correlation between species abundance and biomass (Elbrecht & Leese, 2015) and between species abundance and relative or absolute sequence read abundance (Elbrecht et al., 2017). Our results validate and reinforce the use of sequence abundance information for diversity analyses and interpretation of our DNA metabarcoding data to be used with greater confidence.

### Different taxonomic and numerical resolutions on diversity, community structure and composition

In contrast to our hypothesis that different taxonomic levels and numerical resolution from both methods lead to varying multivariate community patterns, we report similar spatial patterns of community structure and composition based on the outcomes of ordination techniques (NMDS ordinations and Procrustes tests). Moreover, high Procrustes coefficients were found for comparisons between different numerical resolutions rather than the different taxonomic levels. This contrasts with a previous report by Pires et al. (2021), where comparisons between different taxonomic resolutions have higher coefficients than numerical resolution from invertebrate communities. Still, we detected relatively high and significant congruence between the ordinations performed for all the taxonomic and numerical resolutions of both methods and indicate a satisfactory surrogacy between the two methods and their varying identification levels. Our results are similar to previous studies that reported strong congruence between different taxonomic levels from freshwater macroinvertebrate samples (e.g., Brito et al., 2018; Godoy et al., 2019; Caradima, Reichert & Schuwirth, 2020). Similar community distribution patterns may be due to parallel responses to environmental gradients and similar dispersal abilities among taxonomically or evolutionary close taxa, leading to a strong community congruence at different taxonomic resolutions (Heino & Soininen, 2007; Landeiro et al., 2012).

We observed significant differences in the community structure and composition at the river scale between the dam-impacted Trinity River and its pristine tributaries. This observation corroborates with previous studies that comprehensively studied and reported the negative impacts of reservoir and dam construction on macroinvertebrate diversity and community composition (e.g., Monaghan et al., 2005; Serrana et al., 2018; Wang et al., 2020). Interestingly, although all datasets have significant positive correlations based on Procrustes tests, the percentage of the variance explained (*R*^*2*^) by the river scale groups at the family level was higher than at finer taxonomic resolutions (i.e., genus, species, and ASV-levels) for both absolute abundance and presence/absence data. This implies that even if different taxonomic and numerical resolutions are correlated and commonly showed significant differences at the river scale, family-level datasets provided relatively higher variance explained by the dam-impact, and the percentage of variance explained decreased as the taxonomic resolution became finer, i.e., order from family, to genus, to species, and to the ASV-levels.

Likewise, Godoy et al. (2019) reported that family-level identification reduced the unexplained portion of variance compared with genus-level identification of aquatic insect community samples. Family-level identifications have been proposed as surrogates in freshwater macroinvertebrate assemblages for ecological monitoring and conservation management (Chessman, Williams & Besley, 2007; Brito et al., 2018). Here, we reported that DNA metabarcoding and morphology-based family-level data of both numerical resolutions are enough to reveal the damming impacts on the differences of the macroinvertebrate community structure and composition.

Furthermore, significant differences in the community structure and composition were also reported for most of the morphology and DNA metabarcoding datasets of varying taxonomic and numerical resolutions at the reach scale, except for both the ASV levels and the morphology-based family-level (presence/absence) datasets. The non-significant difference from the ASV dataset for both absolute abundance and presence/absence composition indicates that the increased taxonomic resolution, i.e., intraspecific diversity (haplotypes), was not robust enough for community-based analyses and resulted in the insignificant difference of macroinvertebrate community composition at the reach scale. Moreover, the observed non-significant difference on the morphology-based family level presence/absence dataset can be accounted for by the turnover of rare families at the reach scale. The analysis based on presence/absence tends to increase the influence on the spatial distributions of rare taxa in the dataset (Bailey, Norris & Reynoldson, 2001). However, for this dataset, 25 out of the 31 identified families were shared by the gravel bars and non-gravel bar reaches, with only three unique families on each category. Among-reach scale differences in community composition were not significantly observed because most of the families were distributed across these reaches.

For each of the alpha diversity metrics, we found significant and strong correlations between most of the numerical resolution datasets of the Chao1 richness and Shannon diversity index, with a majority of correlations exceeding the 0.7 threshold of congruence suggested by Heino (2010). This suggests a relatively congruent estimation of these alpha diversity metrics at varying taxonomic levels, regardless of numerical resolution. On the other hand, the Berger-Parker dominance and rare taxa abundance indices showed mostly non-significant, even negative correlation between most of the DNA metabarcoding and morphology-based datasets because the former method introduces more taxa undetected from the latter approach, specifically rare taxa, thus, the different patterns in dominance and rarity indices (Gibson et al., 2015).

### Community and environmental relationships at different taxonomic and numerical resolutions

We found the significant influence of environmental variables on community composition for all the different taxonomic and numerical resolution datasets of both methods, but each dataset showed varying sets of associated physicochemical parameters. In particular, the DNA metabarcoding genus, species, and ASV-level datasets showed non-significant associations with some of the environmental variables that were significant for the morphology-based family and genus-level datasets. Some environmental parameters, e.g., electric conductivity, dissolved oxygen and ash-free dry mass, influenced both family-level datasets, while pH only showed significant associations with the DNA metabarcoding species and ASV-level datasets. Finer resolution in taxonomic information introduces more unique or rare taxa which could be attributed to more sensitive response to certain environmental gradients, while coarser taxonomic resolutions detect a lower number of rare taxa (Emilson et al., 2017) which may lead to biased, and a more limited range of environmental factors detected for their influence on the macroinvertebrate communities.

Previous DNA metabarcoding-based macroinvertebrate biomonitoring studies documented stronger discriminatory power due to the finer taxonomic resolution of the method (e.g., Gibson et al., 2015). However, our environmental association analyses presented coarser taxonomic resolution of DNA metabarcoding was more congruent with the morphology-based datasets. Our observation corresponds with former studies. For example, Emilson et al. (2017) reported that increased statistical power from finer resolution to detect the influence of environmental gradients could be study-specific. Bailey, Norris, and Reynoldson’s (2001) reported that additional information from finer taxonomic resolutions may introduce undesirable ecological noise (i.e., a more comprehensive description of the benthic macroinvertebrate community) unless the impacts of environmental disturbances are more evident with finer-level identification than coarser taxonomic resolution. Other studies also reported that using coarser taxonomic resolution showed small or inconspicuous loss of information compared to species-level identifications for the estimation of the relationships between communities and environmental conditions (e.g., Heino & Soininen, 2007; de Oliveira et al., 2020), with some even reporting lower efficiency of finer taxonomic resolution for the environmental associations due to the ecological noise (e.g., Heino, 2014).

For the relationship between richness and the environmental variables, we present that the explanatory power of most of the environmental variables was relatively higher for the morphology-based datasets, but both showed a decrease in explanatory power from finer to coarser taxonomic resolution. Alpha diversity measures such as richness can potentially be strongly underestimated at coarser taxonomic resolution (Maurer, 2000; Mueller, Pander & Geist, 2013), specifically for morphology-based identifications (Emilson et al., 2017), which could be the reason for this decrease in explanatory power. Heino (2014) observed similar patterns for richness-environment relationships where the explanatory power of environmental variables decreased from fine to coarser taxonomic resolution.

### Implications for DNA metabarcoding-based biomonitoring and restoration assessment

The primary goal of ecological assessments or biomonitoring is to precisely detect biological response to environmental change and impact based on community structure and composition rather than a comprehensive and detailed description of the community (Bailey, Norris & Reynoldson, 2001). The identification of a coarser taxonomic resolution could be advantageous for DNA metabarcoding-based applications in situations where the lack of taxonomic information, e.g., poor reference database, might severely affect the quality of biological assessments for all major taxonomic groups. Nonetheless, it should be noted that the convenience of using coarser taxonomic levels should be based on the redundant and in parallel environmental responses of the multiple finer taxa, e.g., species or populations, in the same coarser taxonomic group (Balmford, Lyon & Lang, 2000).

Different ecological patterns and processes are scale-dependent (Bracken et al., 2017; de Oliveira et al., 2020). In this study, most of the datasets revealed similar patterns of community composition and structure to differentiate both at river and reach scales but failed to differentiate at river × reach scale. We expected to observe difference in the community composition of the restored gravel bars along the dam-influenced river against the gravel bars in its tributaries since most of the gravel bars in the dam-influenced river were constructed via fluvial deposition of locally added sediments or by mechanical construction of gravel islands and bars, whereas the tributaries assessed have naturally created gravel bars (Gaeuman, 2014). However, the different datasets did not present significant differentiation of the community structure and composition.

To differentiate the gravel bar communities in the dam-influenced river against the gravel bars in the tributaries, and the communities from non-gravel bar sites, we performed indicator taxa analysis and successfully determined genera and species-level taxa in distinguishing the four groups within the river × reach scale (i.e., DAM+GB+, DAM+GB-, DAM-GB+, and DAM-GB-). Both methods identified *Simulium* as a significant indicator genera of the dam-impacted Trinity River’s gravel bar sites. However, only the DNA metabarcoding dataset showed significant adjusted *p*-values after a false discovery rate (FDR) estimation proving the method’s robustness compared to the morphology-based identification.

*Simulium* is a genus of blackflies from the family Simuliidae (Diptera). DNA metabarcoding identified nine *Simulium* species, i.e., *S. vittatum, S. vittatum* complex, *S. saxoxum, S. defoliarti, S. tuberosum* complex, *S. canadense, S. bracteatum, S. arcticum* complex, and *S. annulitarse*, whereas morphological identifications were only made up to the genus-level. Additionally, the LEfSe analysis on the DNA metabarcoding dataset identified *S. vittatum* alongside *Dicosmoecus gilvipes, Hesperoperla pacifica*, and *Sialis californica* as indicator species of the dam-impacted reaches with gravel bars. Blackfly larvae are dominant suspension feeders in most lotic ecosystems (Zhang et al., 1998) and are considered efficient and opportunistic colonizers and filterers with an important association with suspended particles and predators. They inhabit fast-flowing environments that assure high amounts of transported materials (Lock, Adriaens, & Goethals, 2014). Simuliid communities reflect environmental conditions and have been used as indicators of ecological restoration or degradation, given that their composition is closely related to physicochemical variables and morphological characteristics of running water environments (Kazanci, 2006). The recovery of *Simulium* spp. and these other taxa indicate the restoration effect of gravel bar constructions in the dam-fragmented Trinity river.

## CONCLUSION

Information on the influence of different taxonomic and numerical resolutions of benthic macroinvertebrate community composition and structure on their responses to environmental disturbance or management, e.g., river restoration, may relieve certain limitations or challenges in the aim for finer taxonomic resolution, i.e., species or haplotype-level assignment from DNA metabarcoding-based assessments. Here we compared the performance of different taxonomic and numerical resolutions of DNA metabarcoding with consequent comparison to traditional morphology-based identification and how it affects assessment outcomes on the response of benthic macroinvertebrate communities to the restoration or construction of gravel bars conducted in the dam-impacted Trinity River, with the non-dam influenced tributaries serving as the reference sites. DNA metabarcoding detected 93% of the taxa identified with morphological identification and provided finer taxonomic resolution to the species and haplotype (ASV) levels. We also reported significant correlations between morphological sample abundance, biomass, and DNA metabarcoding read abundance, validating and reinforcing the reliability of using sequence abundance for downstream diversity analyses. This also supports the potential of DNA metabarcoding’s application for quantitative analyses. Moreover, we observed a relatively high and significant congruence in macroinvertebrate community structure and composition of different taxonomic and numerical resolutions of both methods, indicating a satisfactory surrogacy between the two approaches and their varying identification levels and data transformation. Although the community-environmental association were significant for all datasets but showed varying significant associations against the physicochemical parameters, our observations still imply that coarser taxonomic resolution could be advantageous for DNA metabarcoding-based applications in situations where the lack of taxonomic information, e.g., poor reference database, might severely affect the quality of biological assessments.

## Acknowledgments

This work was supported by the Japan Society for the Promotion of Science (JSPS) Grant-in-Aid for Scientific Research (Grant No. 17H01666, 19K21996, and 19H02276). We are grateful to the members of the Disaster Prevention Research Institute, Kyoto University, and Dr. David Gaeuman of the Trinity River Restoration Program for their assistance during the field survey. We thank Dr. Naohito Tokunaga of the Division of Analytical Bio-Medicine for his assistance in performing high-throughput sequencing on the Illumina MiSeq platform of the Advanced Research Support Center (ADRES), Ehime University.

## Data availability statement

The raw sequence data were deposited into the National Center for Biotechnology Information (NCBI) Sequence Read Archive (SRR12620160). The interactive Krona charts, ASV sequences, morphological and metabarcoding data matrices, taxonomy and sample metadata, and the scripts for the bioinformatics analyses were deposited in the Figshare data repository at https://doi.org/10.6084/m9.figshare.15035412.v1 (Serrana et al., 2021).

## Author contributions

K.W., B.L. and J.M.S. conceptualized and designed the study; J.M.S., B.L., Y.T., and T.S. conducted fieldwork; B.L. performed the morphological identification and laboratory work; J.M.S. performed bioinformatics analyses and data interpretation. The manuscript was drafted by J.M.S. with writing inputs and revisions from all coauthors.

## Supplementary information

The supplementary data to this article is provided as a supplemental file (DOCX).

