## Supporting Information for "Implications of taxonomic and numerical resolution on DNA metabarcoding-based inference of benthic macroinvertebrate responses to river restoration"

Running Title: Taxonomic and numerical resolution on metabarcoding

### Authors and Affiliations

Joeselle M. Serrana[1,2], Bin Li[1], Tetsuya Sumi[3], Yasuhiro Takemon[3], and Kozo Watanabe[1,2]

[1]Center for Marine Environmental Studies, Ehime University, Bunkyo-cho 3, Matsuyama, Ehime 790-8577, Japan

[2]Graduate School of Science and Engineering, Ehime University, Bunkyo-cho 3, Matsuyama, Ehime 790-8577, Japan

[3]Disaster Prevention Research Institute, Kyoto University, Gokasho, Uji, Kyoto, 611-0011, Japan

### Correspondence

Prof. Kozo Watanabe, Ph.D.; Molecular Ecology and Health (MEcoH) Laboratory, Engineering Building No.2, Ehime University, Bunkyo-cho 3, Matsuyama, Ehime 790-8577, Japan;

### Supplementary Table

**TABLE S1** Alpha diversity estimates at different scales, and taxonomic resolution for both morphological (“Morp”) and metabarcoding-based (“Meta”) methods.

| Taxonomic Rank | | | River[a] | | | | | Reach[a] | | | | | River × Reach[b] | | | | |
|---|---|---|---|---|---|---|---|---|---|---|---|---|---|---|---|---|---|
| | | | *Richness* | *Shannon* | *Pielou's* | *Berger-Parker* | *Rarity* | *Richness* | *Shannon* | *Pielou's* | *Berger-Parker* | *Rarity* | *Richness* | *Shannon* | *Pielou's* | *Berger-Parker* | *Rarity* |
| Absolute Abundance | **Morp** | Family | 0.03 | 0.06 | 0.50 | 0.49 | 0.07 | 0.44 | 0.74 | 0.19 | 0.20 | 0.53 | 0.12 | 0.14 | 0.37 | 0.25 | 0.16 |
| | | Genus | 0.10 | 0.12 | 0.16 | 0.10 | 0.06 | 0.99 | 0.84 | 0.74 | 0.95 | 0.70 | 0.30 | 0.33 | 0.51 | 0.29 | 0.14 |
| | **Meta** | Family | 0.02 | 0.91 | 0.08 | 0.03 | 0.67 | 0.88 | 0.63 | 0.70 | 0.87 | 0.45 | 0.04 | 0.82 | 0.20 | 0.10 | 0.95 |
| | | Genus | 0.05 | 0.20 | 0.93 | 0.65 | 0.76 | 0.96 | 0.58 | 0.45 | 0.96 | 0.48 | 0.13 | 0.41 | 0.71 | 0.88 | 0.90 |
| | | Species | 0.03 | 0.10 | 0.53 | 0.24 | 0.08 | 0.89 | 0.45 | 0.27 | 0.40 | 0.70 | 0.03 | 0.33 | 0.70 | 0.65 | 0.49 |
| | | ASV | 0.25 | 0.93 | 0.61 | 0.94 | 0.79 | 0.48 | 0.15 | 0.03 | 0.25 | 0.88 | 0.26 | 0.49 | 0.08 | 0.59 | 0.20 |
| Presence/Absence | **Morp** | Family | 0.04 | 0.02 | NA | 0.01 | 0.05 | 0.73 | 0.38 | NA | 0.25 | 0.08 | 0.12 | 0.14 | 0.92 | 0.15 | **0.01** |
| | | Genus | 0.14 | 0.17 | NA | 0.17 | 0.03 | 0.92 | 0.92 | NA | 0.96 | 0.64 | 0.30 | 0.48 | 0.73 | 0.58 | 0.13 |
| | **Meta** | Family | 0.04 | 0.01 | NA | **0.00** | 0.12 | 0.87 | 0.68 | NA | 0.64 | 0.19 | 0.03 | 0.02 | 0.42 | 0.02 | 0.17 |
| | | Genus | 0.09 | 0.03 | NA | 0.02 | **0.00** | 0.98 | 0.97 | NA | 0.95 | 0.51 | 0.13 | 0.12 | 0.76 | 0.14 | 0.02 |
| | | Species | 0.04 | 0.02 | NA | 0.02 | 0.58 | 0.89 | 0.91 | NA | 0.88 | 0.80 | 0.03 | 0.04 | 0.82 | 0.06 | 0.66 |
| | | ASV | 0.23 | 0.28 | NA | 0.31 | 0.66 | 0.38 | 0.66 | NA | 0.84 | 0.12 | 0.17 | 0.41 | 0.67 | 0.54 | 0.54 |

Significant differences (t-test[a]; ANOVA[b]) are indicated in highlight (<0.05) and bold print (<0.01). "NA" indicates that data are essentially constant for statistical test.

**TABLE S2** Environmental data collected for each sampling sites. "EC" stands for electric conductivity; "DO" for dissolved oxygen; "$NH_4$-N" for ammonium nitrogen; "AFDM" for ash-free dry mass; "TSS" for total suspended solids; and "$NO_3$-N" for nitrate nitrogen.

| **Sampling Site** | | **pH** | **EC** (µS/cm) | **DO** (mg/L) | **$NH_4$-N** (mg/L) | **AFDM** ($mg/m^3$) | **TSS** (mg/L) | **$NO_3$-N** (mg/L) |
|---|---|---|---|---|---|---|---|---|
| DAM+GB+ | **B1DZ** | 6.90 | 82 | 10.30 | 0.03 | 200 | 0.90 | 0.04 |
| | **B1UZ** | 6.80 | 81 | 5.94 | 0.02 | 400 | 1.10 | 0.04 |
| | **B2DZ** | 7.40 | 80 | 5.99 | 0.03 | 400 | 1.30 | 0.04 |
| | **B2UZ** | 7.00 | 82 | 14.98 | 0.02 | 400 | 2.60 | 0.06 |
| | **B3DZ** | 7.10 | 85 | 5.55 | 0.02 | 1,500 | 19.50 | 0.06 |
| | **B3UZ** | 7.50 | 82 | 5.35 | 0.01 | 1,200 | 7.60 | 0.04 |
| | **B4DZ** | 7.50 | 86 | 4.71 | 0.02 | 200 | 0.90 | 0.06 |
| | **B4UZ** | 7.10 | 127 | 0.76 | 0.02 | 800 | 22.60 | 0.09 |
| | **B5DZ** | 7.90 | 85 | 6.12 | 0.01 | 400 | 0.80 | 0.06 |
| | **B5UZ** | 7.50 | 85 | 5.14 | 0.01 | 1,800 | 10.40 | 0.05 |
| | **B6DZ** | 7.10 | 85 | 7.15 | 0.01 | 200 | 1.10 | 0.06 |
| | **B6UZ** | 7.40 | 81 | 11.65 | 0.01 | 400 | 4.30 | 0.06 |
| DAM-GB+ | **B7DZ** | 7.80 | 119 | 6.05 | 0.01 | 52,400 | 1.80 | 0.00 |
| | **B7UZ** | 7.60 | 118 | 4.74 | 0.01 | 82,100 | 2.40 | 0.00 |
| | **B8DZ** | 7.40 | 67 | 4.71 | 0.01 | 12,800 | 1.00 | 0.05 |
| | **B8UZ** | 7.80 | 65 | 4.25 | 0.01 | 25,200 | 11.40 | 0.05 |
| DAM-GB- | **F1US** | 7.70 | 117 | 5.86 | 0.02 | 12,400 | 1.10 | 0.05 |
| | **F1DS** | 7.50 | 117 | 6.69 | 0.06 | 9,890 | 0.90 | 0.00 |
| | **F2US** | 7.80 | 65 | 9.47 | 0.01 | 13,100 | 1.00 | 0.00 |
| | **F2DS** | 7.80 | 64 | 5.34 | 0.02 | 9,290 | 0.90 | 0.00 |
| DAM+GB- | **F3US** | 7.40 | 94 | 13.16 | 0.04 | 615 | 0.80 | 0.00 |
| | **F3DS** | 7.60 | 92 | 12.17 | 0.01 | 575 | 0.90 | 0.00 |
| | **F4US** | 7.50 | 92 | 14.82 | 0.03 | 33 | 0.50 | 0.00 |
| | **F4DS** | 7.70 | 96 | 13.01 | 0.01 | 968 | 0.70 | 0.00 |

Sampling site abbreviations: "DAM-GB-" stands for Reference Non-gravel bar reach; "DAM-GB+" stands for Reference Gravel Bars; "DAM+GB-" stands for Trinity Non-gravel bar reach; "DAM+GB+" stands for Trinity Gravel Bars; "DZ" represents the down-welling and "UZ" up-welling zones of a gravel bar (B); "US" represents the up-stream and "DS" down-stream points of collection for the non-gravel bar reaches of the rivers (F) (approx. 20-m length).

**TABLE S3** PERMDISP2 analysis to test the effect of the scales on the physicochemical parameters. “EC” stands for electric conductivity; “DO” for dissolved oxygen; “NH4-N” for ammonium nitrogen; “AFDM” for ash-free dry mass; “TSS” for total suspended solids; and “NO3-N” for nitrate nitrogen.

| **Scale** | | **pH** | **EC** (uS/cm) | **DO** (mg/L) | **$NH_4$-N** (mg/L) | **AFDM** ($mg/m^3$) | **TSS** (mg/L) | **$NO_3$-N** (mg/L) |
|---|---|---|---|---|---|---|---|---|
| **River** | Df | 1 | 1 | 1 | 1 | 1 | 1 | 1 |
| | Sum Sq | 0.00 | 0.00 | 0.02 | 0.00 | 0.00 | 0.05 | 0.08 |
| | Mean Sq | 0.00 | 0.00 | 0.02 | 0.00 | 0.00 | 0.05 | 0.08 |
| | F value | 2.29 | 46.14 | 2.08 | 0.03 | 1.40 | 1.10 | 1.28 |
| | Pr(>F) | 0.14 | 0.00 *** | 0.16 | 0.88 | 0.25 | 0.31 | 0.28 |
| **Reach** | Df | 1 | 1 | 1 | 1 | 1 | 1 | 1 |
| | Sum Sq | 0.00 | 0.00 | 0.00 | 0.03 | 0.00 | 0.17 | 0.02 |
| | Mean Sq | 0.00 | 0.00 | 0.00 | 0.03 | 0.00 | 0.17 | 0.02 |
| | F value | 3.86 | 0.41 | 0.11 | 2.11 | 0.18 | 14.05 | 0.36 |
| | Pr(>F) | 0.06 . | 0.53 | 0.74 | 0.16 | 0.67 | 0.00 ** | 0.56 |
| **River × Reach** | Df | 3 | 3 | 3 | 3 | 3 | 3 | 2 |
| | Sum Sq | 0.00 | 0.00 | 0.03 | 0.08 | 0.01 | 0.21 | 0.16 |
| | Mean Sq | 0.00 | 0.00 | 0.01 | 0.03 | 0.00 | 0.07 | 0.08 |
| | F value | 2.57 | 14.75 | 0.84 | 1.02 | 1.00 | 3.98 | 1.36 |
| | Pr(>F) | 0.08 . | 0.00 *** | 0.49 | 0.40 | 0.41 | 0.02 * | 0.29 |

Note: “River” (2 groups: Trinity River – dam-influenced, Reference River – non-dam-influenced); "Reach" (2 groups: gravel bars, free-flowing reach). Significance code: ‘***’ associated with a variable at p < 0.001, ‘**’ at p < 0.01, ‘*’ at p < 0.05 and ‘.’ at p < 0.1.

**TABLE S4** List of genera identified as indicators at the “River × Reach” scale using multivariate pattern analysis (Association function: IndVal.g).

| **Genus** | **DAM-GB-** | **DAM-GB+** | **DAM+GB-** | **DAM+GB+** | **index** | **stat** | **p.value** | **fdr.p.value** |
|---|---|---|---|---|---|---|---|---|
| Morphology-based | | | | | | | | |
| *Rhyacophila* | 1 | 1 | 1 | 0 | 11 | 0.92 | 0.009 | 0.1800 |
| *Simulium* | 0 | 0 | 0 | 1 | 4 | 0.92 | 0.022 | 0.2170 |
| *Nemoura* | 0 | 0 | 1 | 1 | 10 | 0.82 | 0.048 | 0.3227 |
| DNA Metabarcoding | | | | | | | | |
| ***Wormaldia*** | **1** | **1** | **0** | **0** | **5** | **0.99** | **0.000** | **0.0075** |
| ***Hydropsyche*** | **1** | **1** | **0** | **0** | **5** | **0.96** | **0.003** | **0.0868** |
| ***Antocha*** | **1** | **0** | **0** | **0** | **1** | **0.87** | **0.006** | **0.0868** |
| ***Attenella*** | **1** | **0** | **0** | **0** | **1** | **0.87** | **0.006** | **0.0868** |
| ***Haemaphysalis*** | **0** | **1** | **0** | **0** | **2** | **0.81** | **0.006** | **0.0868** |
| ***Ordobrevia*** | **1** | **1** | **0** | **0** | **5** | **0.86** | **0.007** | **0.0868** |
| ***Simulium*** | **0** | **0** | **0** | **1** | **4** | **0.95** | **0.008** | **0.0868** |
| *Diphetor* | 1 | 1 | 0 | 0 | 5 | 0.77 | 0.026 | 0.2006 |
| *Lara* | 1 | 1 | 0 | 0 | 5 | 0.71 | 0.027 | 0.2006 |
| *Optioservus* | 1 | 0 | 0 | 0 | 1 | 0.82 | 0.028 | 0.2006 |
| *Sweltsa* | 0 | 0 | 1 | 0 | 3 | 0.82 | 0.032 | 0.2006 |
| *Epeorus* | 0 | 1 | 1 | 1 | 14 | 0.94 | 0.032 | 0.2006 |
| *Thienemannimyia* | 0 | 1 | 0 | 0 | 2 | 0.68 | 0.042 | 0.2394 |

Only the indicator taxa with significance value <0.05 are shown in the table. Indicator taxa with false detection rate adjusted p-value <0.10 are highlighted in bold. “DAM-GB-” stands for Reference Non-gravel bar reach; “DAM-GB+” stands for Reference Gravel Bars; “DAM+GB-” stands for Trinity Non-gravel bar reach; “DAM+GB+” stands for Trinity Gravel Bars; “fdr.p.value” stands for the false detection rate adjusted p-value. Columns with the 0 and 1s indicate which site groups were included in the combination preferred by the taxa. Index indicates the index of the site group combination. The remaining two columns are the association statistic, the p-value of the permutational test.

### Supplementary Figures

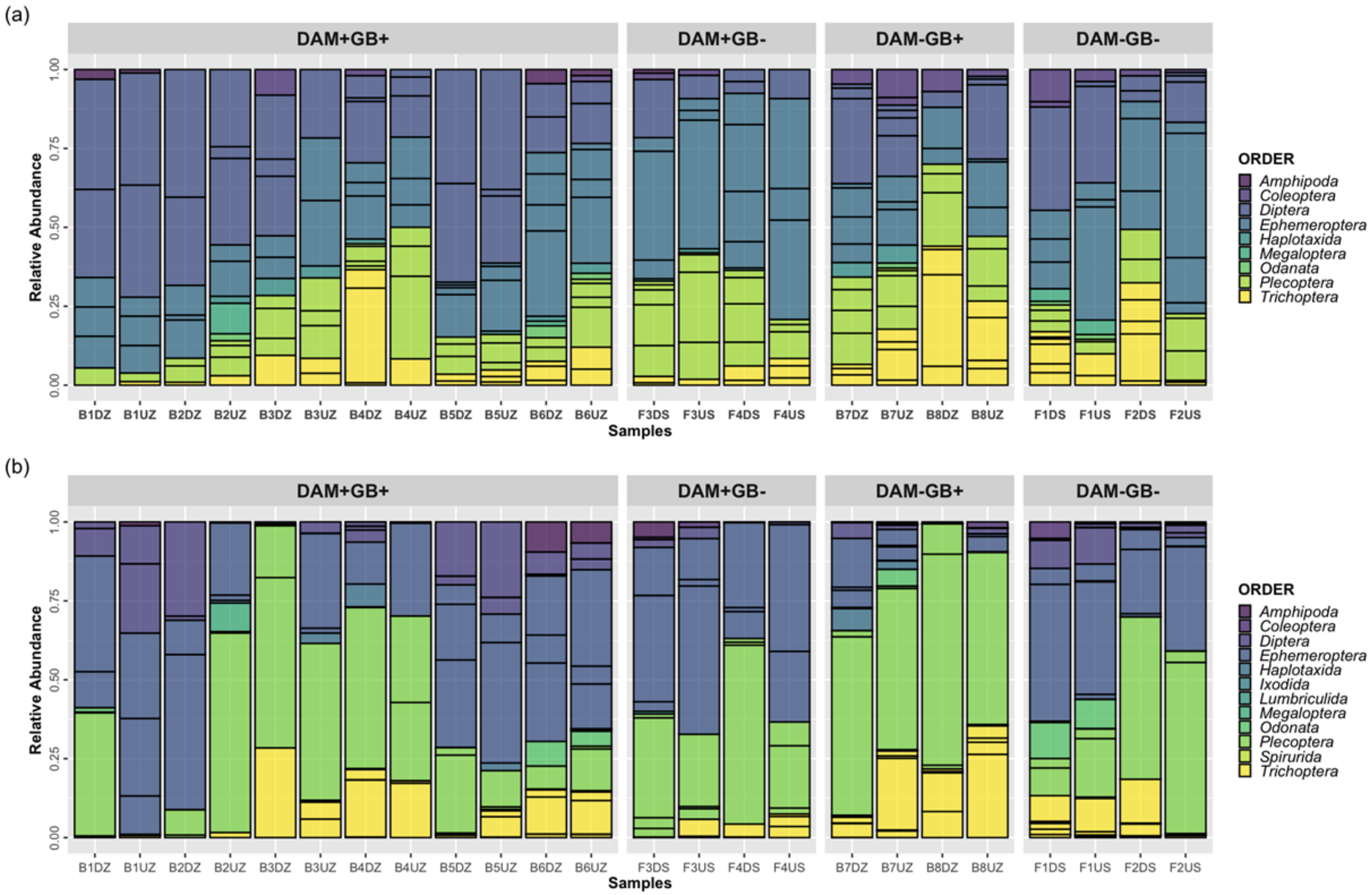


**FIGURE S1** Relative abundance of the morphological (a) and metabarcoding (b) datasets at the Order-level.

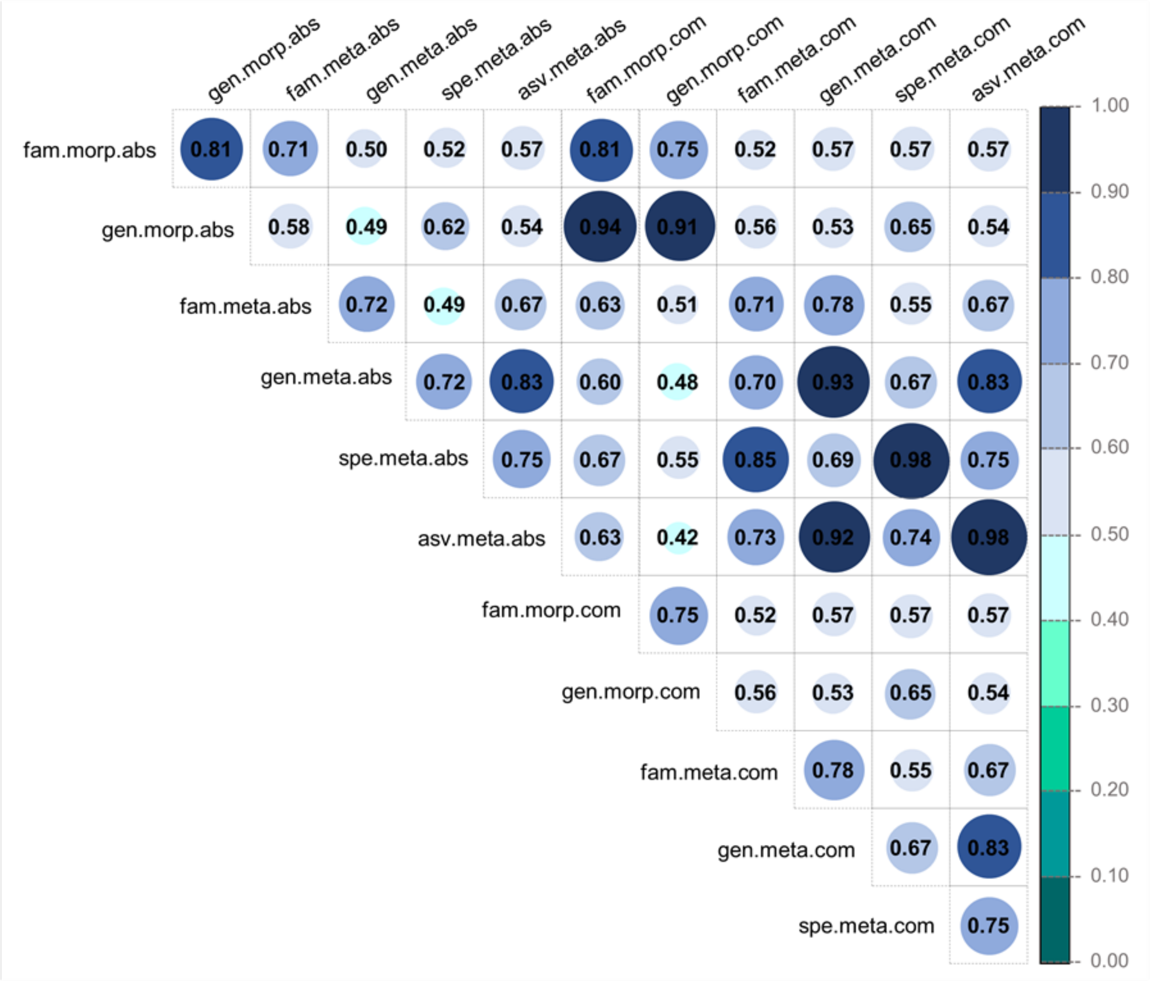


**FIGURE S2** Correlogram of the Procrustes analyses of the dissimilarity matrices of the different taxonomic ranks of the morphology and metabarcoding datasets. “morp” indicates morphologically identified; “meta” for DNA metabarcoding; “gen” for genus-level identification; “fam” for family-level; “spe” for species–level; “asv” for the amplicon sequence variant (ASV) level dataset; “abs” for absolute abundance; “com” for presence/absence data. Values in the circles represent the correlation in a symmetric Procrustes rotation. Correlation boxes with Xs are not significant at 0.05.

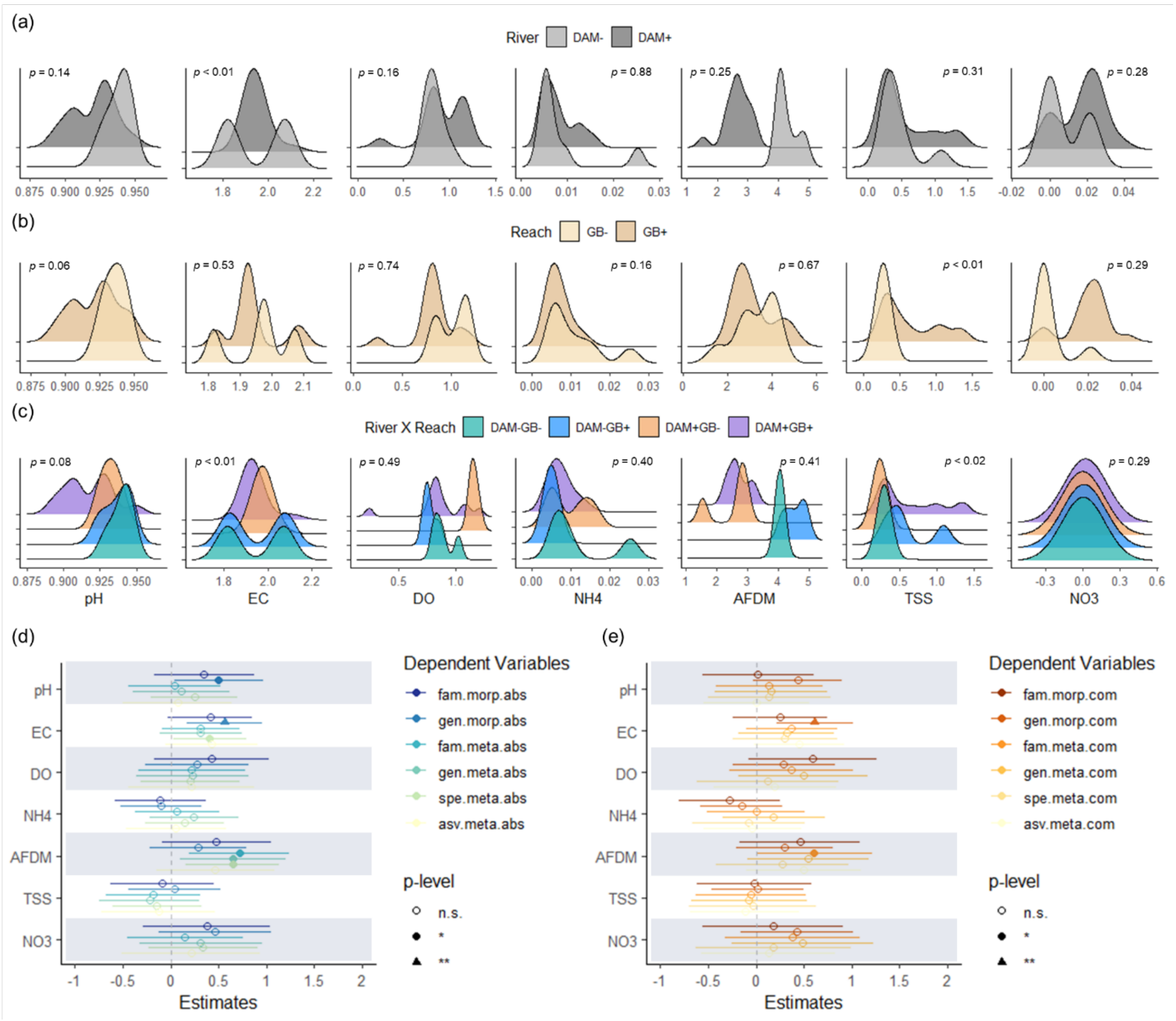


**FIGURE S3** Distribution plots of the environmental variables (log-transformed) per scale: River (a), Reach (b) River × Reach (c). "EC" stands for electric conductivity; "DO" for dissolved oxygen; "$NH_4$-N" for ammonium nitrogen; "AFDM" for ash-free dry mass; "TSS" for total suspended solids; and "$NO_3$-N" for nitrate nitrogen. *p*-values from the PERMDISP2 analysis to test the effect of the scale on the physicochemical parameters (Supplementary Table S3). Forest plot of the meta-regression analysis of the environmental variables with Chao1 richness as dependent variable for the absolute abundance (d) and presence/absence (e) data. "morp" indicates morphologically identified; "meta" for DNA metabarcoding; "gen" for genus-level identification; "fam" for family-level; "spe" for species–level; "asv" for the amplicon sequence variant (ASV) level dataset; "abs" for absolute abundance; "com" for presence/absence data. "*" statistical significant at $p < 0.05$, "**" statistical significant at $p < 0.01$.

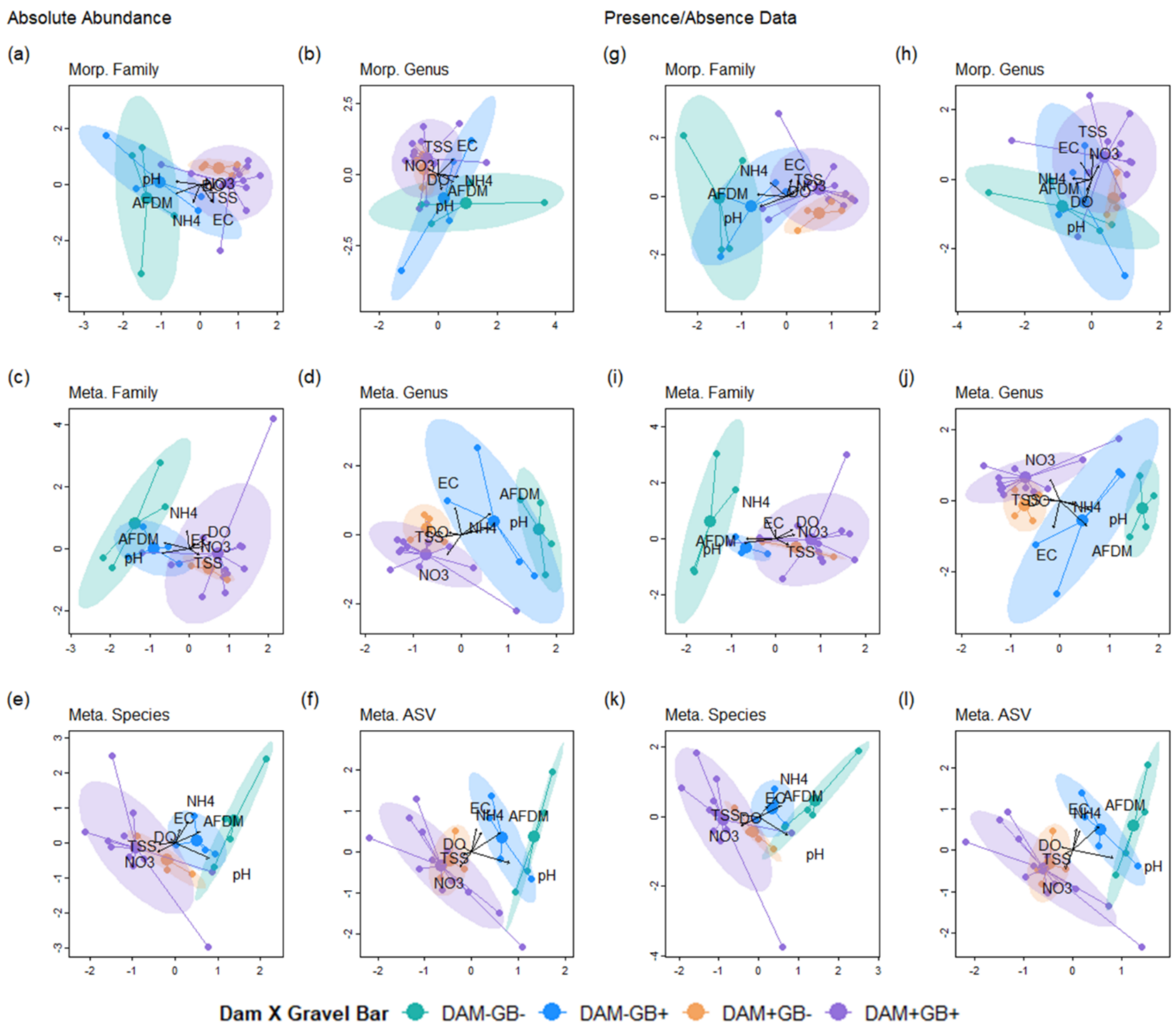


**FIGURE S4** Canonical correspondence analysis (CCA) biplot based on Bray-Curtis dissimilarity for absolute abundance (a-f) and presence/absence (g-l) data of the macroinvertebrates detected by morphological ("Morp.") and metabarcoding-based ("Meta.") methods.

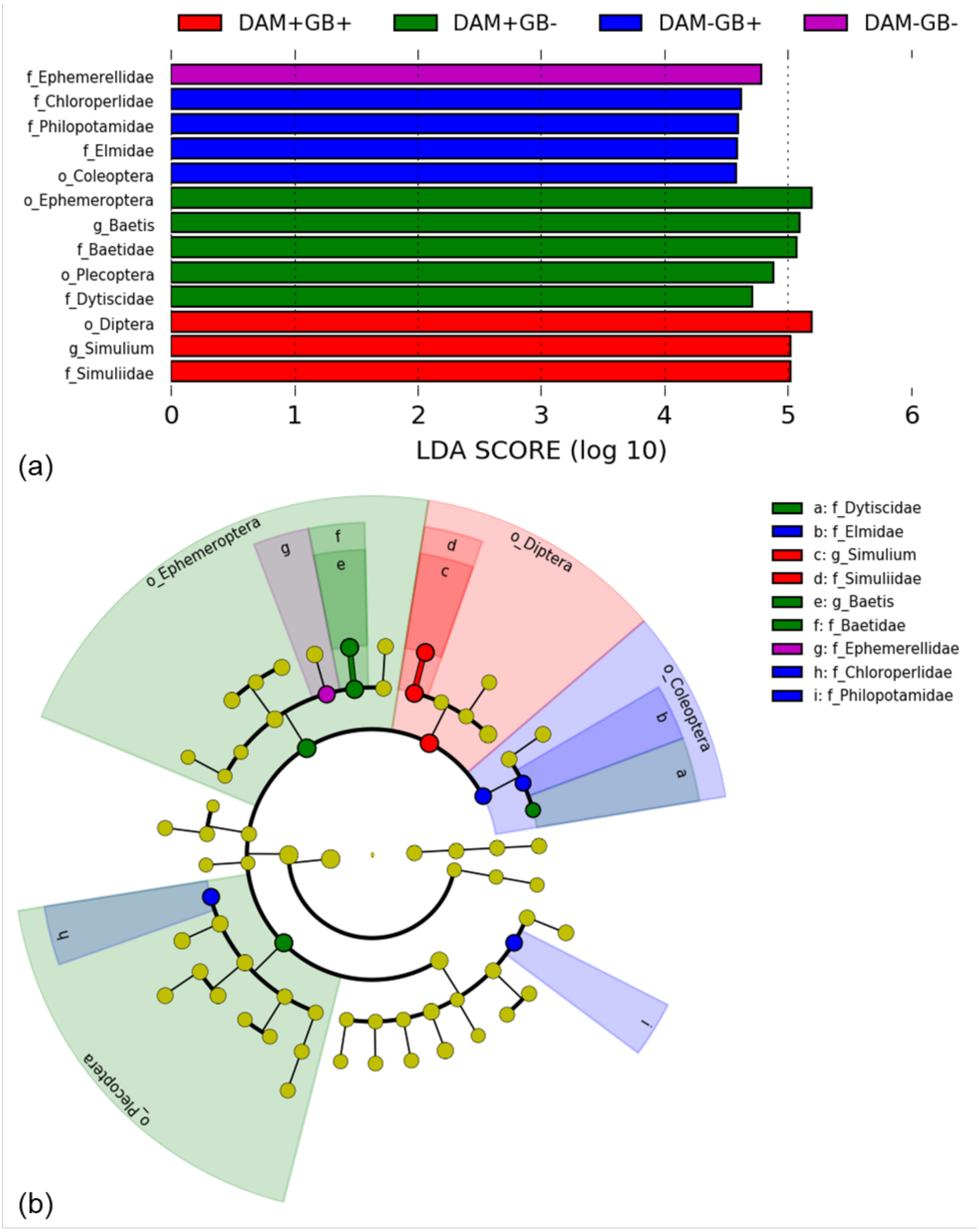


**FIGURE S5** Linear Discriminant Analysis (LDA) Effect Size (LEfSe) plot of indicator taxa for the morphologically-identified dataset. Identified indicator taxa grouped by River × Reach scale and ranked by effect size (a). The threshold for LDA score was > 2.0. Cladogram representing the hierarchical structure of the indicator taxa identified (b).

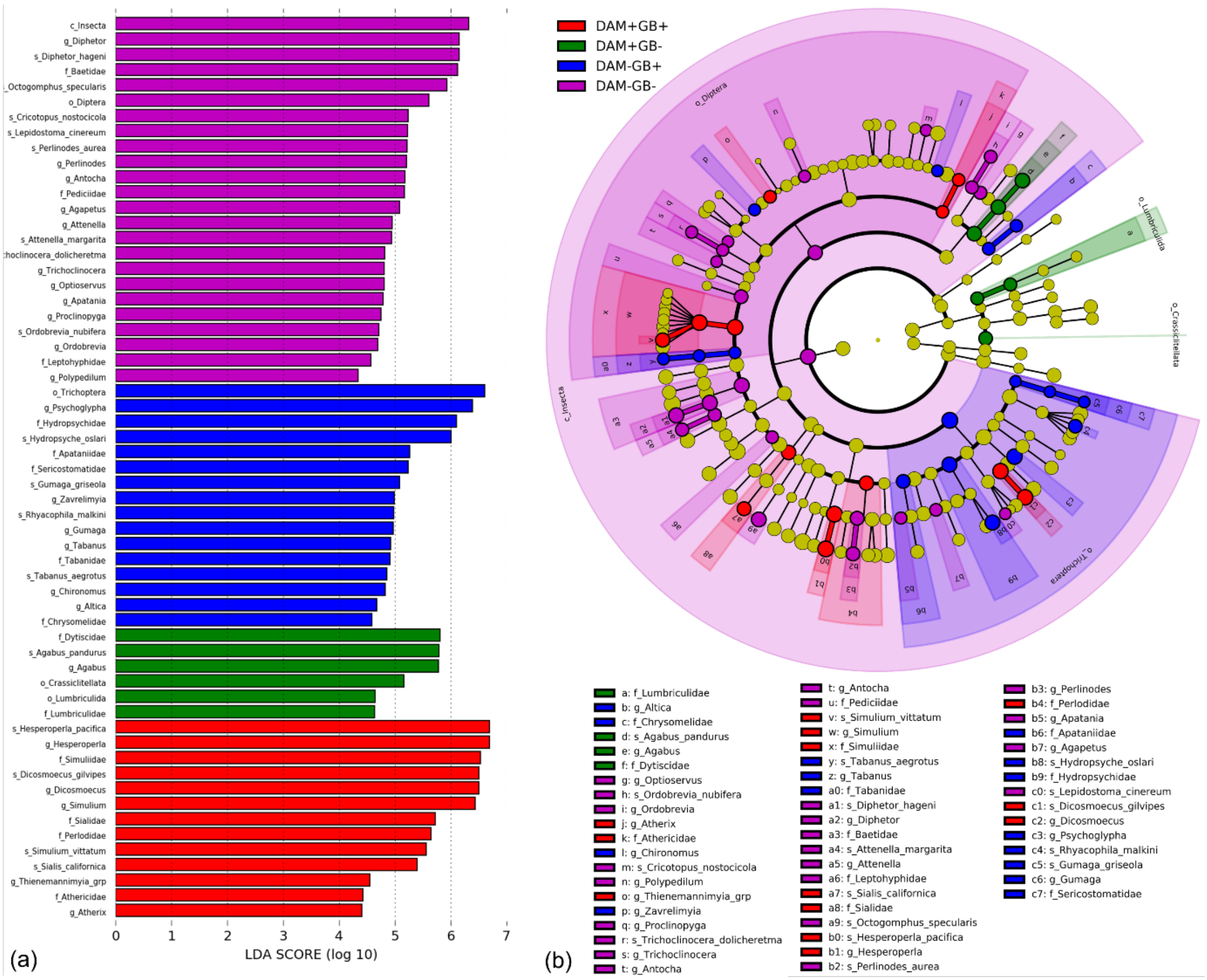


**FIGURE S6** Linear Discriminant Analysis (LDA) Effect Size (LEfSe) plot of indicator taxa for the DNA metabarcoding dataset. Identified indicator taxa grouped by River × Reach scale and ranked by effect size (a). The threshold for LDA score was > 2.0. Cladogram representing the hierarchical structure of the indicator taxa identified (b).
